# Global Profiling of Differentiating Macrophages Identifies Novel Functional Long Non-coding RNAs Regulating Polarization and Innate Immune Responses

**DOI:** 10.1101/2023.04.09.536159

**Authors:** Araceli M. Valverde, Raza A. Naqvi, Afsar R. Naqvi

**Author notes:** Corresponding Author: Afsar Naqvi, Ph.D., Associate Professor, Department of Periodontics, College of Dentistry, Adjunct Associate Professor, Department of Microbiology and Immunology, College of Medicine, Mucosal Immunology Lab Room 561 B, University of Illinois Chicago Chicago, IL 60612.

## Abstract

Macrophages (Mφ) are functionally dynamic immune cells that bridge innate and adaptive immune responses. However, the underlying epigenetic mechanisms that control the macrophage plasticity and innate immune functions are not well-elucidated. Here we performed transcriptome profiling of differentiating M1Mφ and M2Mφ and identified thousands of previously known and novel lncRNAs. We characterized three Mφ-enriched lncRNAs (LRRC75A-As1, GAPLINC and AL139099.5) with novel functions in Mφ differentiation, polarization and innate immunity. Knockdown of LRRC75A-As1, and GAPLINC downregulated Mφ differentiation markers CDw93 and CD68, and skewed macrophage polarization by decreasing M1 markers but had no significant impact on M2 markers. LRRC75A-As1, and GAPLINC RNAi in Mφ attenuated bacterial phagocytosis, antigen processing and inflammatory cytokine secretion supporting their functional role in potentiating innate immune functions. Mechanistically, lncRNA knockdown perturbed the expression of multiple cytoskeleton signaling thereby impairing Mφ migration suggesting their critical role in regulating macrophage polarity and motility. Together, our results show that Mφ acquire a unique repertoire of lncRNAs to shape differentiation, polarization and innate immune functions.

## Introduction

The ability of a healthy immune system to recognize and clear a plethora of antigens rely on the enormous plasticity displayed by the comprising cell types. Monocytes and macrophages (Mφ) are members of the mononuclear phagocyte system (MPS) that constantly patrol the peripheral tissues and are actively recruited to the sites of injury and infection (*1–5*). Infiltrating monocytes, under the influence of tissue microenvironment, can differentiate to Mφ or Dendritic cells (DCs). In tissues, mature Mφ are activated in response to a combination of local stimuli to acquire specialized functional phenotypes *viz*., inflammatory (M1) and reparative (M2) (*6–11*). Equipped with diverse Toll-like receptors (TLRs) these cells of innate arm of immunity recognize and phagocytize antigens, secrete cytokines that activate the adaptive arm of the immune system and perform key roles in wound repair (*1–4*). In contrast, overt immune responses by infiltrated immune cells can contribute to the pathogenesis of numerous diseases including rheumatoid arthritis, periodontitis, etc., (*12–15*). Deciphering the role of endogenous regulatory molecules that control differentiation and function of these multifaceted immune cells may uncover novel mechanisms to control their phenotypes.

Advances in transcriptome sequencing enabled by human genome annotation has revealed pervasive transcriptional activity across the genome (*16–18*). Intriguingly, significantly higher number of newly sequenced transcripts exhibit little or no protein-coding potential. Non-coding transcripts longer than 200 nucleotides (nts) are an emerging class of regulatory RNA defined as long non-coding RNAs (lncRNAs). Unlike protein-coding genes, lncRNAs exhibit high tissue specificity, temporal expression, poor sequence conservation, and relatively lower expression (*16–18*). Moreover, lncRNAs can be found both in nuclear and cytoplasmic compartments, while mRNAs are predominantly present in cytoplasm indicating broader regulatory role of lncRNAs (*17,18*). The ability of lncRNAs to physically interact with DNA, RNA or protein allows them to regulate transcriptional, post-transcriptional as well as translational output of the genome (17-*19*). Not surprisingly, aberrations in the lncRNA expression has been implicated in various diseases including cancers, neurodegenerative disorders, autoimmunity, etc., (*22–25*).

Differentiation of Mφ from monocytes is critical in the activation of immune responses against any pathological threat and subsequent tissue repair and homeostasis (*1–5*). Overt immune activity can be equally detrimental. While numerous lncRNAs are reported in myeloid cells their raison d’être is largely unknown. Considering that approximately 25-40% protein-coding genes have overlapping antisense transcription, the impact of lncRNAs on gene regulation is not to be underestimated (*15–19*). Few reports have demonstrated lncRNA-mediated control of myeloid cell differentiation and function. Huang et al., (2016) selected six differentially expressed lncRNAs in M1 and M2 polarized macrophages including CMPK2, THRIL, TCONS_00019715, ENST00000569328, ENST00000414554 and ENST00000474886 because of their potential roles in modulating the inflammatory response of macrophages. TCONS_00019715 was responsive to M1/M2 switch and its knockdown suppressed M1 markers and upregulated M2 markers suggesting its role in polarization. PACERR, also known as COX-2-lncRNA, controls the inflammatory response of macrophages by regulating COX-2 transcripts (*11*). A recent study has shown that lncRNA PACERR plays critical role in the polarization of tissue-associated macrophages (TAMs). The expression levels of M2 markers CD206, CD163, Arginase-1, TGF-β, and IL-10 were downregulated after PACERR knockdown in TAMs (*26*). Tian et al. demonstrated that lncRNA LINC00662 promotes M2 macrophage phenotype associated with hepatocellular carcinoma (HCC) progression by activating Wnt/β-Catenin signaling favoring tumor growth and metastasis by upregulating WNT3A (*27*). However, the mechanisms through which lncRNAs regulate macrophages innate immune functions remains less explored.

In this study, we comprehensively profiled lncRNA repertoire during the differentiation of monocytes to M1Mφ and M2Mφ using high-throughput RNA sequencing (RNAseq) and characterized their mechanistic role in modulating myeloid cell differentiation and functions. In addition to previously reported lncRNAs, we identified large number of novel lncRNAs sequences that accumulate in specific stages of macrophage differentiation. Our findings provide the most up-to-date expressional dynamics of lncRNA repertoire during macrophage differentiation and functionally characterized three previously unannotated lncRNAs *viz*., LRRC75A-As1, GAPLINC and AL139099.5 in regulating polarization, innate immune activity and migration.

## Results

### Dynamic changes in lncRNA expression profile occur during macrophage polarization

LncRNAs exhibit tissue-, cell-, and developmental stage-specific expression and are differently expressed under physiological or pathological conditions (*16–18*). We therefore performed time-kinetics of lncRNA profiling in monocyte-derived M1 and M2 Mφ to identify lncRNA repertoire expressed in differentiating macrophages. We isolated total RNA from monocytes (Mon: n=3), and differentiating M1Mφ and M2Mφ at 18h (M1; n=4/M2; n=4), day 3 (M1; n=5/M2; n=4), day 5 (M1; n=5/M2; n=4) and day 7 (M1; n=4/M2; n=4) and examined lncRNA profiles by RNA-seq. A total of 270 million (monocytes), 363 million (M1Mφ-18h), 512 million (M1Mφ-day 3), 509 million (M1Mφ-day 5), 464 million (M1Mφ-day7), 422 million (M2Mφ-18h), 365 million (M2Mφ-day 3), 409 million (M2Mφ-day 5) and 426 million (M2Mφ-day 7) raw reads were obtained by RNA-seq (**Table 1**). After quality control, approximately, 192, 291, 416, 379, 335, 343, 298, 301 and 284 million of clean reads were obtained for monocytes, M1Mφ-18h, M1Mφ-day 3, M1Mφ-day 5, M1Mφ-day 7, M2Mφ-18h, M2Mφ-day 3, M2Mφ-day 5, and M2Mφ-day 7, respectively (**Table 1**). In all the samples ∼75-79% reads uniquely mapped to the human genome, except M2Mφ-day 5 where we obtained 66% mapped reads (**Table S1**). Sequences that aligned with the reported lncRNAs were considered as known lncRNAs and these transcripts were classified as class j, i, o, and x (**Fig. 1 and Data S1**). Approximately, each class of lncRNA group were represented by 3% (j class), 70% (I class), 4% (o class), and 3% (x class) lncRNA transcripts (**Fig. 1**). Another properties, such as transcript lengths, exon number and open reading frame (ORFS) of coding genes and lncRNAs in every condition were also analyzed (**Fig. S1 A-D**). In compliance with MIAME Guidelines, we deposited the RNA-seq datasets to Gene Expression Omnibus public database under the Accession Number **GSE192642**.

**Fig. 1.**
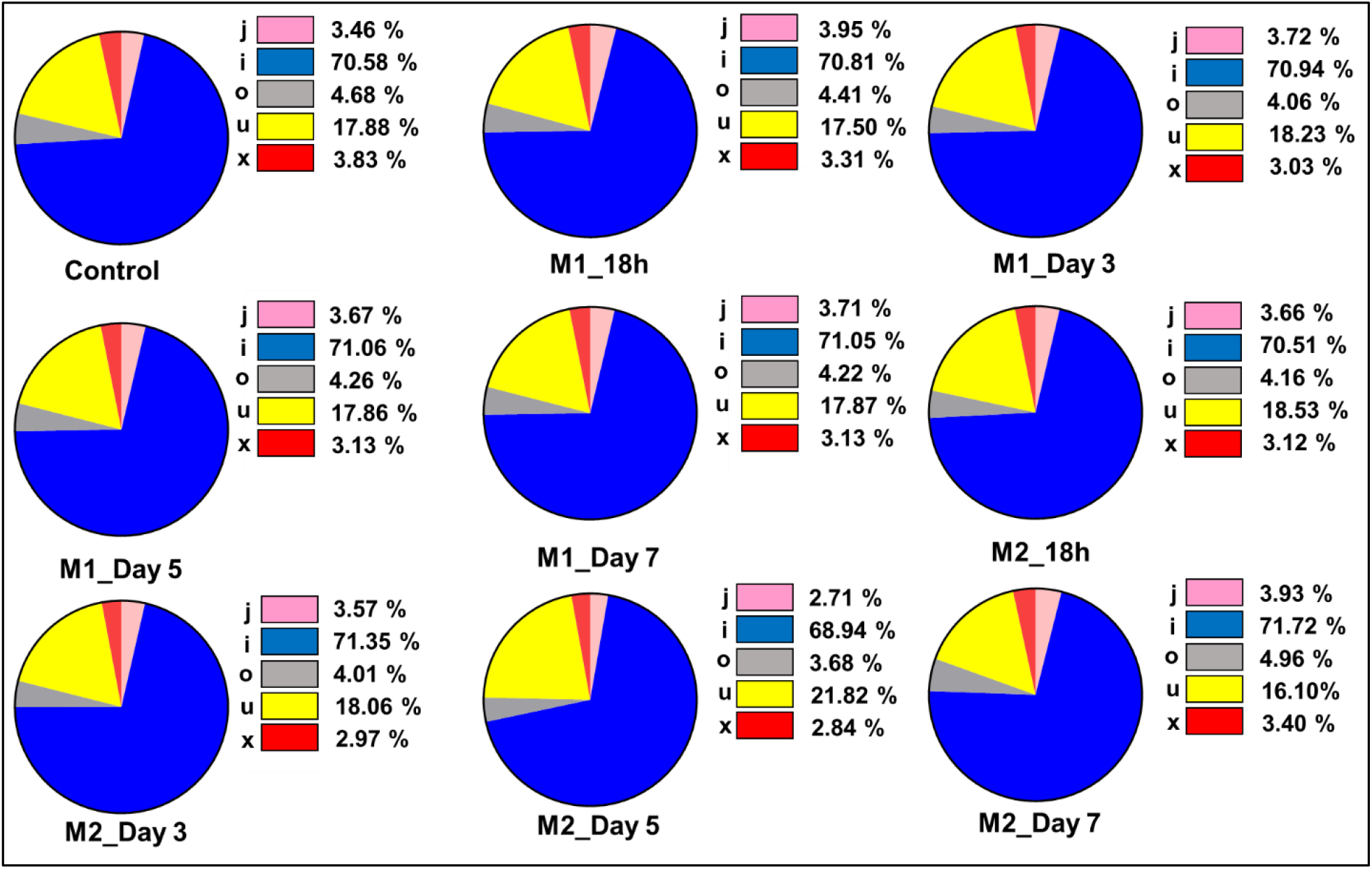
Different lncRNA classes identified in differentiating macrophages. LncRNAs were subdivided into five categories according to their class code generated by StringTie. (i) a transfrag falling entirely within a reference intron (intronic); (j) potentially novel isoform or fragment at least one splice junction is shared with a reference transcript; (o) generic exonic overlap with a reference transcript; (u) unknown, intergenic transcript (intergenic); (x) Exonic overlap with ref erence on the opposite strand (antisense). Approximately, each class of lncRNAs group were represented by 3%, (j class) 70% (I class), 4% (o class), 17% (u class) and 3 % (x class).

**Table 1.** Summary of quality control, statistics of reads and sequences reads mapping to genome.

| Sample | Raw Data | Valid reads | Mapped reads | Unique Mapped reads |
| --- | --- | --- | --- | --- |
| Mon_C | 92531180 | 78316480 | 72338565(92.37%) | 58842681(75.13%) |
| Mon_6 | 81572852 | 66451384 | 62088269(93.43%) | 51374923(77.31%) |
| Mon_7 | 96627304 | 48036478 | 43827794(91.24%) | 36213502(75.39%) |
| M1_18h_1 | 76711890 | 58954182 | 53108875(90.09%) | 39818218(67.54%) |
| M1_18h_2 | 106594234 | 88570634 | 82829990(93.52%) | 67062065(75.72%) |
| M1_18h_3 | 75373348 | 62740726 | 58969269(93.99%) | 48055893(76.59%) |
| M1_18h_4 | 105253564 | 81595936 | 76244615(93.44%) | 61113684(74.90%) |
| M1_Day3_1 | 105861648 | 89010170 | 84537952(94.98%) | 68595066(77.06%) |
| M1_Day3_2 | 100586208 | 82604734 | 77840881(94.23%) | 63984953(77.46%) |
| M1_Day3_3 | 92730726 | 74569970 | 70117444(94.03%) | 56434578(75.68%) |
| M1_Day3_4 | 99500128 | 77519952 | 73107243(94.31%) | 61373055(79.17%) |
| M1_Day3_5 | 113713356 | 92811358 | 86942271(93.68%) | 67734123(72.98%) |
| M1_Day5_1 | 92916138 | 72943162 | 69501973(95.28%) | 58054013(79.59%) |
| M1_Day5_2 | 118448628 | 98807924 | 94063905(95.20%) | 76276898(77.20%) |
| M1_Day5_3 | 83787072 | 65418736 | 61776124(94.43%) | 51147186(78.18%) |
| M1_Day5_4 | 101740162 | 58710800 | 53868690(91.75%) | 43483460(74.06%) |
| M1_Day5_5 | 112950108 | 83288656 | 79572352(95.54%) | 64755965(77.75%) |
| M1_Day7_1 | 87774676 | 72193228 | 68317239(94.63%) | 54897039(76.04%) |
| M1_Day7_2 | 109007820 | 82545880 | 76910569(93.17%) | 60008045(72.70%) |
| M1_Day7_3 | 104642568 | 77938600 | 72485395(93.00%) | 59535860(76.39%) |
| M1_Day7_4 | 162907558 | 103029476 | 95366073(92.56%) | 78534151(76.22%) |
| M2_18h_1 | 122746974 | 101841330 | 94259785(92.56%) | 76939191(75.55%) |
| M2_18h_2 | 98838350 | 78507450 | 70973713(90.40%) | 55652156(70.89%) |
| M2_18h_3 | 99270216 | 79123226 | 73863238(93.35%) | 58763449(74.27%) |
| M2_18h_4 | 101275056 | 83843146 | 78978088(94.20%) | 64431661(76.85%) |
| M2_Day3_1 | 94132538 | 78870856 | 75057569(95.17%) | 63162554(80.08%) |
| M2_Day3_2 | 85895838 | 71672454 | 67729212(94.50%) | 56649293(79.04%) |
| M2_Day3_3 | 85874334 | 70699718 | 67198571(95.05%) | 55959974(79.15%) |
| M2_Day3_4 | 99577974 | 77054454 | 72150571(93.64%) | 60428202(78.42%) |
| M2_Day5_1 | 104344564 | 71592858 | 68304248(95.41%) | 57713357(80.61%) |
| M2_Day5_2 | 131206178 | 107041042 | 101690368(95.00%) | 86658041(80.96%) |
| M2_Day5_3 | 101919572 | 84982940 | 81015568(95.33%) | 68822222(80.98%) |
| M2_Day5_4 | 71553066 | 37613438 | 10300884(27.39%) | 8231557(21.88%) |
| M2_Day7_1 | 94729636 | 76387320 | 71959105(94.20%) | 58288785(76.31%) |
| M2_Day7_2 | 94507236 | 73805298 | 69004752(93.50%) | 57427800(77.81%) |
| M2_Day7_3 | 114857722 | 63230520 | 60078019(95.01%) | 52460632(82.97%) |
| M2_Day7_4 | 122142704 | 71491636 | 62302467(87.15%) | 52054423(72.81%) |

Next, we examined the differentially expressed lncRNAs in M1Mφ and M2Mφ at 18h, day 3, 5 and 7 and compared with monocytes as control group (**Fig. 2A**). Based on the fold change (a cut off between -1.25 and 1.25) and *p* value (<0.05), 4683 lncRNAs were differentially expressed in M1Mφ at 18h (1555 upregulated and 3128 downregulated), 7649 lncRNAs were differentially expressed in M1Mφ at day 3 (1435 upregulated and 6214 downregulated), 1361 lncRNAs were differentially expressed in M1Mφ at day 5 (597 upregulated and 764 downregulated) and 747 lncRNAs were differentially expressed in M1Mφ at day 7 (442 upregulated and 305 downregulated) (**Fig. 2A**). In M2Mφ, we identified 5458 (1266 upregulated and 4192 downregulated), 6673 (1283 upregulated and 5390 downregulated), 1556 (1542 upregulated and 14 downregulated), and 1816 (1524 upregulated and 292 downregulated) differentially expressed lncRNAs at 18 h, day 3, 5 and 7, respectively (**Fig. 2A**). It can be noted that both M1 and M2Mφ show a peak in differentially expressed lncRNAs at day 3, which is a key time-point in the lineage commitment towards M1/M2 phenotype.

**Fig. 2.**
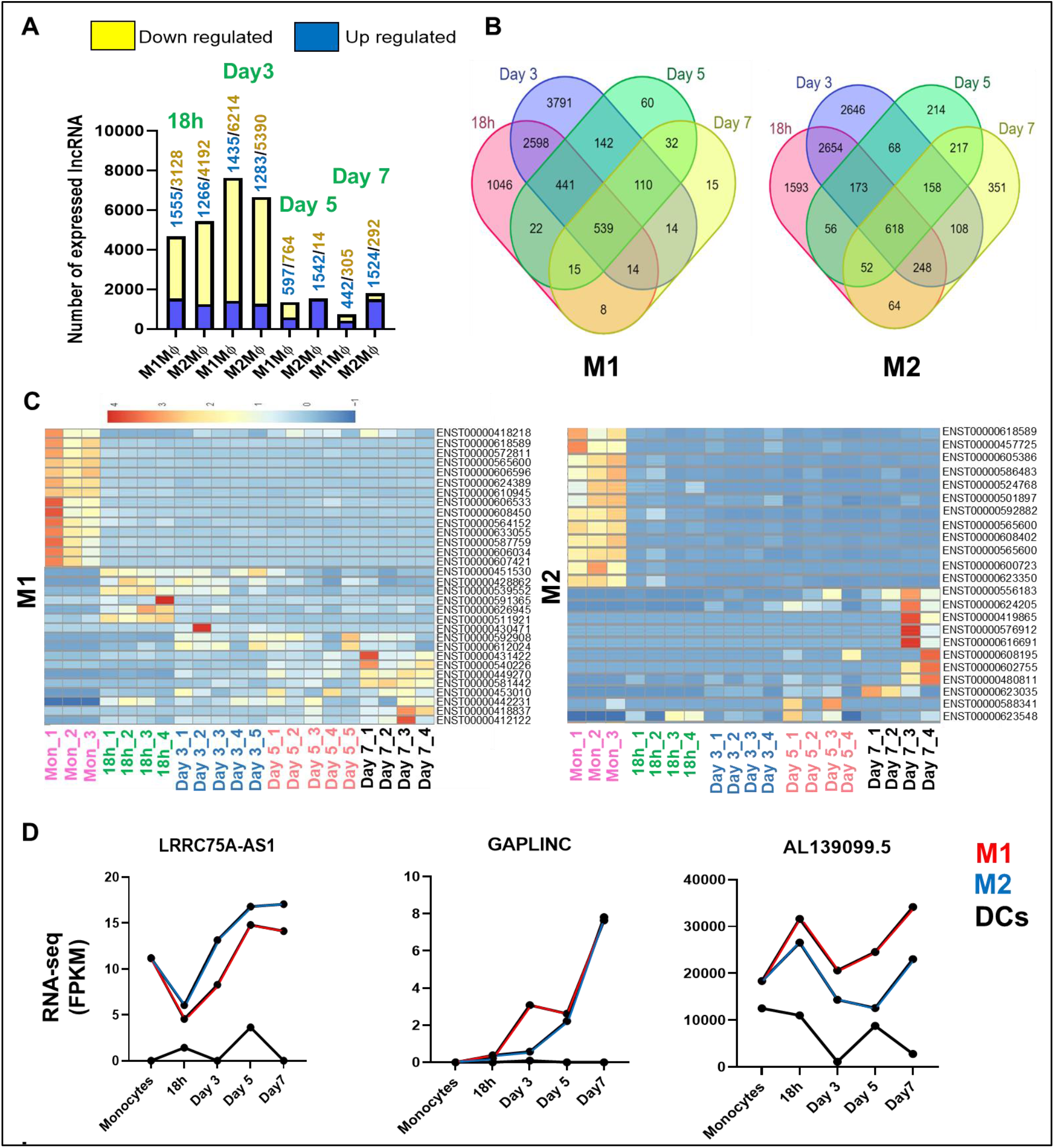
M1 and M2 polarized macrophages express a unique subset of lncRNAs. **A**) LncRNA expression profiles were examined in M1Mφ and M2Mφ at 18h, Day 3, 5 and 7. The number of differentially upregulated and downregulated lncRNAs in each condition were identified based on the fold change (a cut off between -1.25 and 1.25) and p ˂ 0.05 (multiple ANOVA). **B**) 4-set Venn chart showing the number of unique and common lncRNAs expressed in M1Mφ and M2Mφ at 18h, day 3, 5 and 7. **C)** Heatmap showing expression profiles of upregulated and downregulated lncRNAs in M1Mφ and M2Mφ compared to monocytes. **D)** Expression of LRRC75A-AS1, GAPLINC and AL139099.5 in monocytes, M1Mφ, M2Mφ and DC at 18h, day 3, 5 and 7 by calculating FPKM (Total_exon_fragments/mapped_reads(millions)×exon_length(kB). **E)** LRRC75A-AS1, GAPLINC and AL139099.5 expression by RT-qPCR in M1 and M2Mφ. To validate RNAseq results, total RNA was isolated from differentiating macrophages from an independent cohort (n=4) and the expression of LRRC75A-AS1, GAPLINC and AL139099.5 was quantified by RT-qPCR. Actin was used as endogenous control. Data are means ± SEM of three independent donors. The Ct values of three replicates were analyzed to calculate fold change using the 2^−ΔΔCt^ method. Student’s *t*-test was conducted to calculate *p*-values. \**p* < 0.05, \*\**p* < 0.01, \*\*\**p* < 0.001.

Next, we compared lncRNA profiles for each time point and identified a large subset of differentially expressed lncRNAs in polarized Mφ suggesting a specific temporal requirement and expression profiles of lncRNAs. Venn diagram shows the unique and common subset of differentially expressed lncRNAs at 18h, day 3, 5 and 7, in M1Mφ and M2Mφ. A total of 1046, 3791, 60 and 15 lncRNAs were specifically expressed in M1Mφ at 18h, day 3, day 5 and day 7 time points, respectively (**Fig. 2B**). In case of M2Mφ, a total of 1593, 2646, 214 and 351 lncRNAs were specifically expressed during macrophage differentiation. This suggests that acquisition of a unique lncRNA repertoire govern phenotype commitment of M1 and M2 Mφ. Heatmaps in **Fig. 2C** show selected differentially expressed (up- and down-regulated) lncRNAs across different time points in M1Mφ and M2Mφ. Individual heatmaps and Venn diagram comparing monocytes with each Mφ differentiation time-point are presented in **Fig. S3**. Overall, time-kinetics of lncRNA profiles in differentiating M1 and M2 Mφ clearly reveal that lncRNAs are responsive to differentiation stimuli, exhibit temporal expression and encode a unique subset of lncRNAs that define each phenotype.

### A large array of novel lncRNAs sequences identified in differentiating M1 and M2 polarized macrophages

After analyzing the known lncRNAs profiles, we examined our datasets for novel lncRNA sequences. To this end, transcripts that overlapped with known mRNAs, known lncRNAs and transcripts shorter than 200 bp were discarded. Then we utilized CPC (Coding Potential Calculator) (*28*) and CNCI (Coding-Non-Coding Index) to predict transcripts with coding potential (**Fig. S2**) (*29*). All transcripts with CPC score <-1 and CNCI score <0 were removed. We identified a subset of novel lncRNA sequences expressed at specific time-point during macrophage differentiation (**Fig. 3**). **Fig. 3 A,B** shows the number of differentially expressed (up- and down-regulated) novel lncRNAs in M1 and M2 Mφ. Based on the fold change (a cut off between -1.25 and 1.25) and *p* value (<0.05), we identified 765 (109 up and 656 down), 1299 (260 up and 1039 down), 813 (262 up and 561 down) and 636 (286 up and 350 down) novel lncRNAs that were differentially expressed in M1Mφ at 18h, day 3, day 5 and day 7, respectively (**Fig. 3A**). In M2Mφ, 929 (248 up and 681 down), 2017 (1142 up and 875 down), 300 (297 up and 3 down) and 303 (250 up and 53 down) novel lncRNAs were differentially expressed at 18h, day 3, day 5 and day 7, respectively (**Fig. 3A**). Similar to known lncRNA analysis, we noticed highest number of differentially expressed lncRNAs at day 3 further suggesting it as a crucial time-point in the lineage commitment. Heatmaps composed of 20 most significantly up- and down-regulated novel lncRNAs in differentiating M1Mφ and M2Mφ are shown in **Fig. 3C**.

**Fig. 3.**
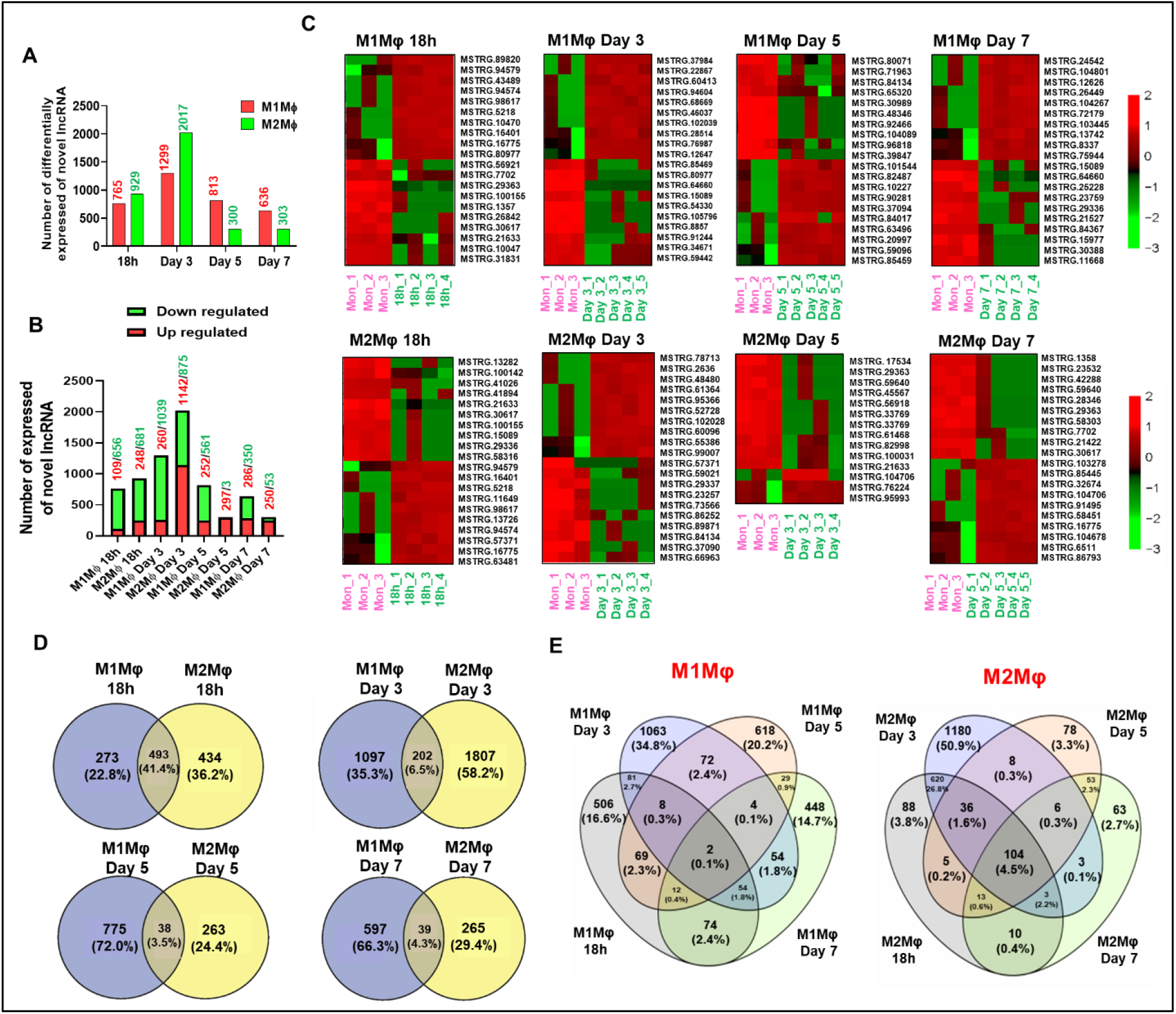
Identification of novel lncRNA sequences during M1 and M2 macrophages polarization. **A**) The total number of differentially expressed novel lncRNAs sequences and, **B**) Number of up- and down-regulated novel lncRNAs identified in M1Mφ and M2Mφ at 18h, day 3, 5 and 7. Sequences were included based on the fold change (a cut off between -1.25 and 1.25) and p ˂ 0.05 in each condition. **C**) Heatmaps showing expression pattern of 20 upregulated and 20 downregulated novel lncRNAs in M1Mφ and M2Mφ compared to monocytes. **D)** Venn diagram showing the number of unique and common novel lncRNAs expressed in M1Mφ and M2Mφ at 18h, day 3, 5 and 7. **D)** 4-set Venn diagram showing the number of specific and common novel lncRNAs expressed in differentiating M1Mφ and M2Mφ.

To identify whether M1Mφ and M2Mφ express unique novel lncRNAs during differentiation, we compared each time point and identified unique and common differentially expressed novel lncRNAs repertoire during M1 and M2 macrophages differentiation (**Fig. 3D**). A total of 1063, 618 and 448 lncRNAs were specifically expressed in monocyte-derived M1Mφ at 18h, day 3, day 5 and day 7, respectively; while in M2Mφ we identified 88, 1180, 78, 63 lncRNAs were specifically expressed during macrophages differentiation. We noticed that a large number of novel lncRNAs exhibit unique expression profiles in M1 and M2 polarized Mφ suggesting temporal transcriptional dynamics. Finally, 4-set flower Venn diagram shows unique and common lncRNAs expressed during M1Mφ and M2Mφ differentiation (**Fig. 3E**). Our results show a large repertoire of previously unreported lncRNA sequences that are temporally expressed during macrophage differentiation.

### Knockdown of candidate lncRNAs impair M1 and M2 differentiation and polarization markers

Next, we investigated whether LRRC75A-AS1, GAPLINC and AL139099.5 play functional role in macrophage differentiation and polarization. Because the candidate lncRNAs were upregulated during differentiation, we examined their impact by RNAi. We designed two siRNAs (A and B) for each lncRNA and transfected them alone or in combination to examine their efficacy in M2Mφ. As shown in the **Fig. 4A**, siRNA-mediated knockdown of LRRC75A-AS1, GAPLINC and AL139099.5 significantly reduced the expression of target transcripts compared to control siRNA. We chose siAL139099.5 A, siGAPLINC B and the combination of siLRRc75A-As1 A+B for the subsequent experiments.

**Fig. 4.**
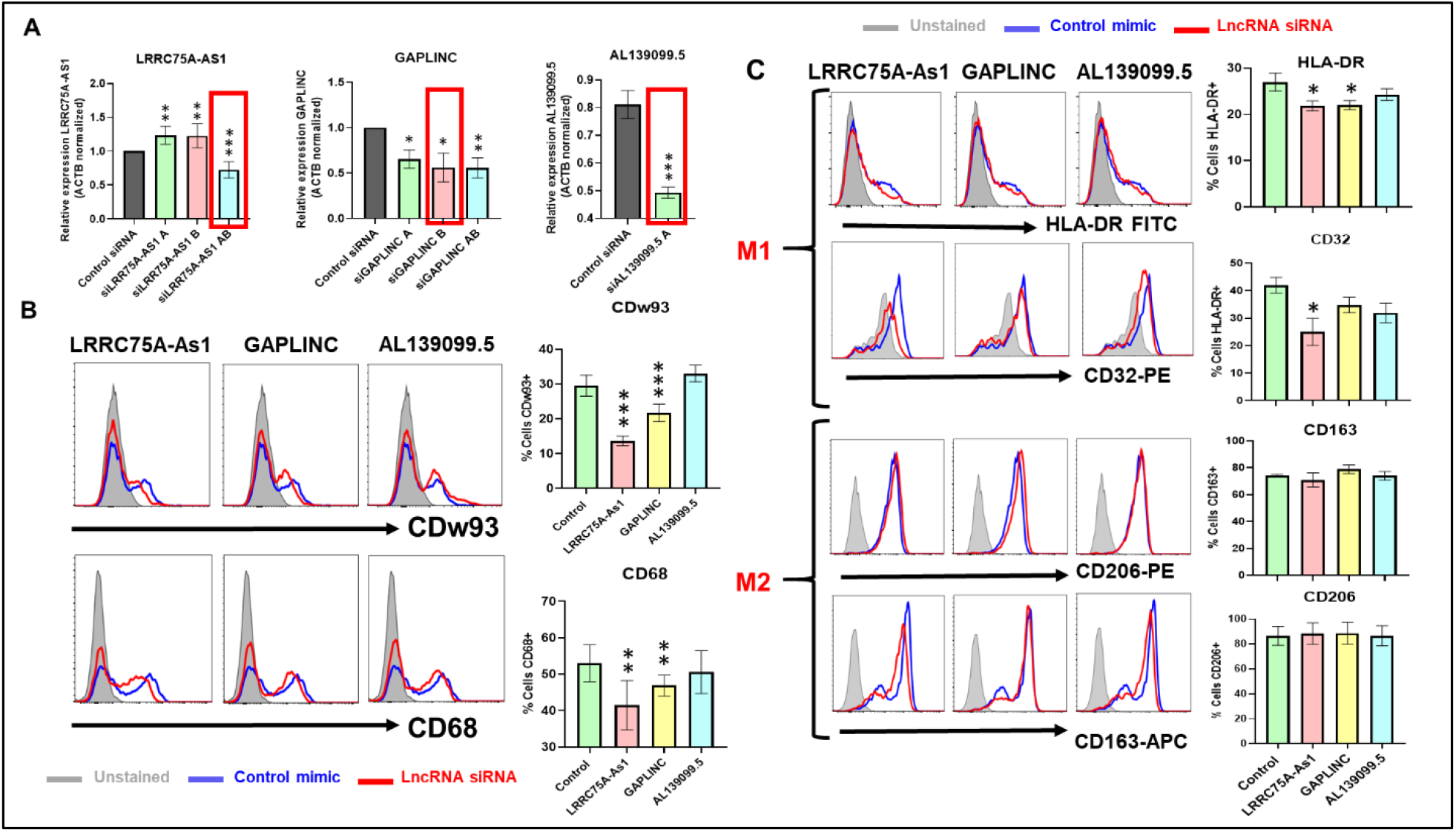
Knockdown of LRRC75A-As1, and GAPLINC impair macrophage differentiation and polarization markers. **A**) Screening of siRNAs targeting candidate lncRNAs by RT-qPCR. M2 macrophages were transfected with two different siRNAs (alone or in combination) targeting LRRC75A-As1, GAPLINC, AL139099.5 or control and the expression of target RNAs were quantified by RT-qPCR. Actin was used as housekeeping gene. Data are means ± SEM of three independent donors. The Ct values of three replicates were analyzed to calculate fold change using the 2^−ΔΔCt^ method. Student’s *t*-test was conducted to calculate *p*-values. \**p* < 0.05, \*\**p* < 0.01, \*\*\**p* < 0.001 **B)** Analysis of differentiation markers in M2 macrophages. M2 macrophages were transfected with selected siRNAs targeting lncRNAs on day 2 and the surface expression of differentiation markers CDw93 and CD68 were analyzed by flow cytometry on day 6. Histograms showing percentage of CDw93 and CD68 positive cells in macrophages. Data are means ± SEM of three independent donors. Student’s *t*-test was conducted to calculate *p*-values. \**p* < 0.05, \*\**p* < 0.01, \*\*\**p* < 0.001. **C)** LRRC75A-As1, and GAPLINC regulate macrophage polarization by suppressing M1 phenotype markers. M1 and M2 macrophages were transfected with candidate lncRNA targeting siRNA or control and M1 and M2 differentiation markers were analyzed by flow cytometry. Histograms showing percentage positive cells of M1 (HLA-DR and CD32) and M2 (CD206 and CD163) polarization markers. Data are means ± SEM of three independent donors. Student’s *t*-test was conducted to calculate *p*-values. \**p* < 0.05, \*\**p* < 0.01.

Day 2 macrophages were transfected with lncRNA-targeting siRNA and the expression of differentiation markers was analyzed by flow cytometry. We observed a significant reduction in the CDw93 and CD68 positive cells was observed in LRRC75A-As1 (13.6±0.4% [CDw93]; 41.4±6.7% [CD68]) and GAPLINC (21.7±2.5% [CDw93]; 46.8±2.9% [CD68] knockdown cells, compared to control (29.6±3.0% [CD6w93]; 52.9±5.1% [CD68]) (**Fig. 4B**). However, no significant changes were noticed for CDw93 (33.1±2.4%) or CD68 (50.5±5.8%) in siAL139099.5 transfected Mφ. These results suggest unique functional activity of candidate lncRNAs in regulating macrophage differentiation.

We next examined if the candidate lncRNA knockdown impair macrophage polarization (**Fig. 4C**). Cells were transfected with lncRNA targeting siRNAs or control at day 3 and the expression of M1 macrophages (HLA-DR and CD32) and M2 macrophages (CD206 and CD163) markers were analyzed on day 6. Flow cytometric analysis show that the expression of M1 macrophages markers (MHCII and CD32) show differential impact of lncRNAs. We observed significant downregulation of HLA-DR+ cells in LRRC75A (21.8±1.0%) and GAPLINC (22±0.9%) but not AL139099.5 (24.3±1.3%) knockdown compared to control (27±1.9%). LRRC75A knockdown reduced (25±4.5% vs control 42±2.8%) CD32+ cells, while no significant changes were observed in GAPLINC (34.8±2.8%) or AL139099.5 (31.9±3.5%) knockdown cells (**Fig. 4C**). On the contrary, none of the lncRNA knockdown showed any impact on M2 markers CD206 and CD163 (**Fig. 4C**; lower panel). Overall, our results show that novel lncRNAs viz., LRRC75A-AS1, GAPLINC are enriched during macrophage lineage commitment, and control differentiation and polarization suggesting their functional requirement during the process.

### LncRNAs LRRC75S-AS1, GAPLINC and AL139099.5 exhibit differential impact on the phagocytosis by polarized macrophages

Phagocytosis is a central role of macrophage activity. Macrophages acquire different phenotype and functions depending on the microenvironment. Both M1 (inflammatory) and M2 (reparative) macrophages perform phagocytosis to remove pathogens and dead cells, respectively, and participate in immune and tissue homeostasis (*30,31*). We therefore examined new biological functions of candidate lncRNAs LRRC75S-AS1, GAPLINC and AL139099.5. M1 and M2Mφ were transiently transfected with the siRNAs targeting LRRC75S-AS1, GAPLINC or AL139099.5 to assess their impact on the phagocytosis of rhodamine labeled *E. coli* bioparticles. Fluorescent microscopy imaging shows attenuated phagocytosis of *E. coli* (red signals) in LRRC75S-AS1, and GAPLINC but not AL139099.5 knockdown cells, compared to control siRNA (**Fig. 5A**). siLRRC75A-As1 showed most significant inhibition of phagocytosis indicating its potent biological activity in macrophages (**Fig. 5A**). Flow cytometric analysis further indicates reduced uptake of *E .coli* in LRRC75S-AS1 (∼35%), and GAPLINC (∼13%) knockdown cells supporting our imaging results (**Fig. 5B**; left panel and **Fig. 5C**; upper panel). Intringuingly, the impact of lncRNA RNAi in M2Mφ was strikingly different than M1Mφ. Compared to scramble, only LRRC75S-AS1 knockdown showed mild attenuation in phagocytosis (∼18%), while no significant changes were observed for GAPLINC or AL139099.5 (**Fig. 5B**; right panel and **Fig. 5C**; lower panel). Overall, these results further support that candidate lncRNAs show a differential functional impact on bacterial phagocytosis in polarized macrophages.

**Fig. 5.**
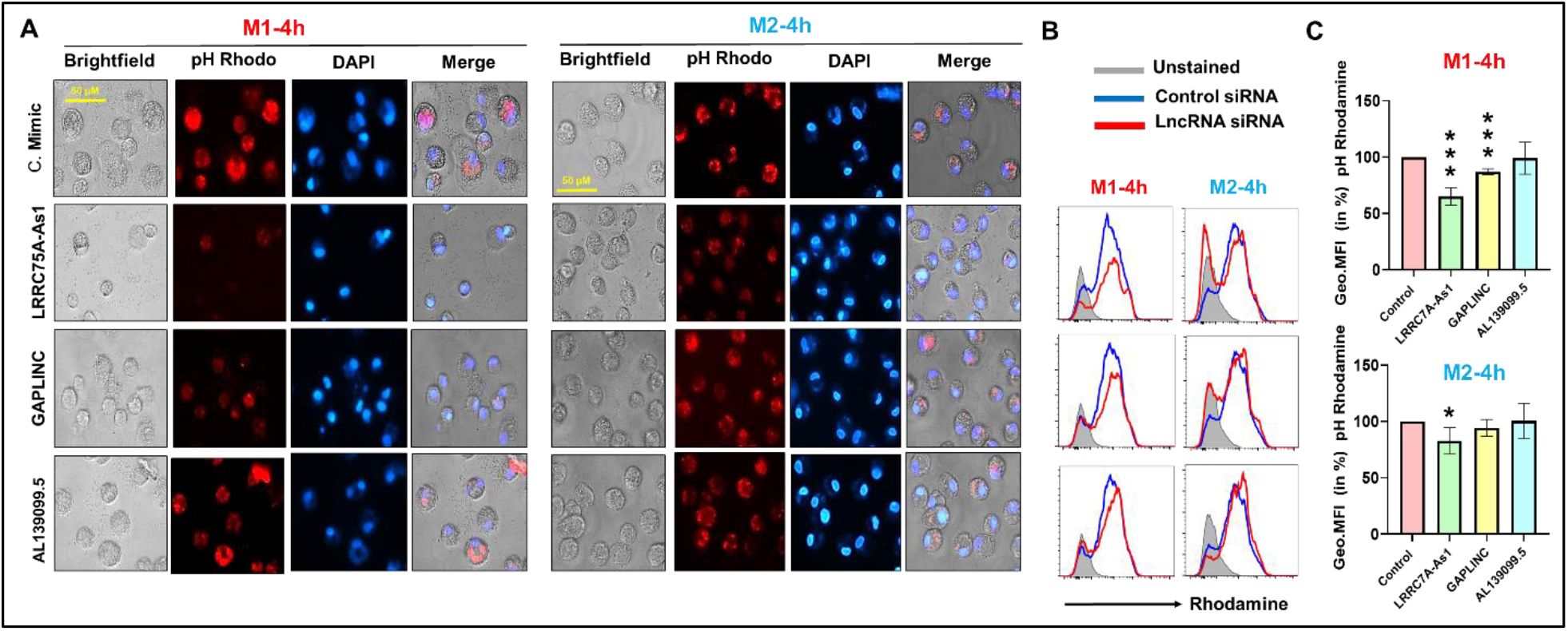
Differential attenuation of phagocytosis in polarized macrophages by lncRNA knockdown. M1 and M2 macrophages were transfected with siRNA targeting LRRC75A-As1, GAPLINC, AL139099.5 or scramble control and phagocytosis was assessed by rhodamine labeled *E. coli* bioparticles. **A)** Representative florescence microscopy images showing bacterial phagocytosis as reflected by rhodamine signal. Final magnification: X40, scale bar corresponds to 100 μm. M1 and M2 macrophages transfected with control or lncRNA targeting siRNAs. **B)** Cells were harvested after 4 h for flow cytometry analysis. Overlay histograms showing attenuation of phagocytosis in LRRC75A-As1, and GAPLINC knockdown cells. **C)** Bar graphs showing percent Geometric MFI in M1 and M2 macrophages. Data are means ± SEM of three independent donors. Student’s *t*-test was conducted to calculate *p*-values. \**p* < 0.05, \*\**p* < 0.01, \*\*\**p* < 0.001.

### Antigen processing in macrophages is regulated by LRRC75S-AS1 and GAPLINC

Phagocytosis of the foreign particles is followed by an effective antigen processing and presentation to T-cells to trigger the adaptive immune response (*32, 33*). However, the role of lncRNAs in this regard remains largely unknown. Next, we analyzed the impact of soluble antigen (Ovalbumin) uptake and processing to demonstrate immune functions of candidate lncRNAs. M1 and M2Mφ transfected with lncRNA-targeting siRNA or control and after 36 h cells were incubated with DQ Ovalbumin to monitor antigen processing. Fluorescent microscopy imaging shows attenuated antigen uptake and processing of Ovalbumin (green signals) in LRRC75S-AS1 and GAPLINC RNAi but no significant changes were observed for AL139099.5 knockdown, compared to control siRNA. However, the most significant inhibition of antigen uptake and processing was observed in LRRC75A-As1 knockdown (**Fig. 6A**). Flow cytometric analysis further showed reduced uptake of Ovalbumin in LRRC75S-AS1 (∼32%) and GAPLINC (∼27%) knockdown M1Mφ supporting our imaging results (**Fig. 6B**; left panel and **Fig. 6C;** upper panel). Similar to phagocytosis assays, the impact of lncRNA RNAi in M2Mφ was strikingly different than M1Mφ. Compared to scramble, only LRRC75S-AS1 knockdown showed mild attenuation in antigen uptake and processing (∼15%), while no significant changes were observed for GAPLINC (**Fig. 6B**; right panel and **Fig. 6C**; lower panel). Overall, these results indicate that our candidate lncRNAs show a differential functional impact of bacterial antigen uptake and processing in polarized macrophages.

**Fig. 6.**
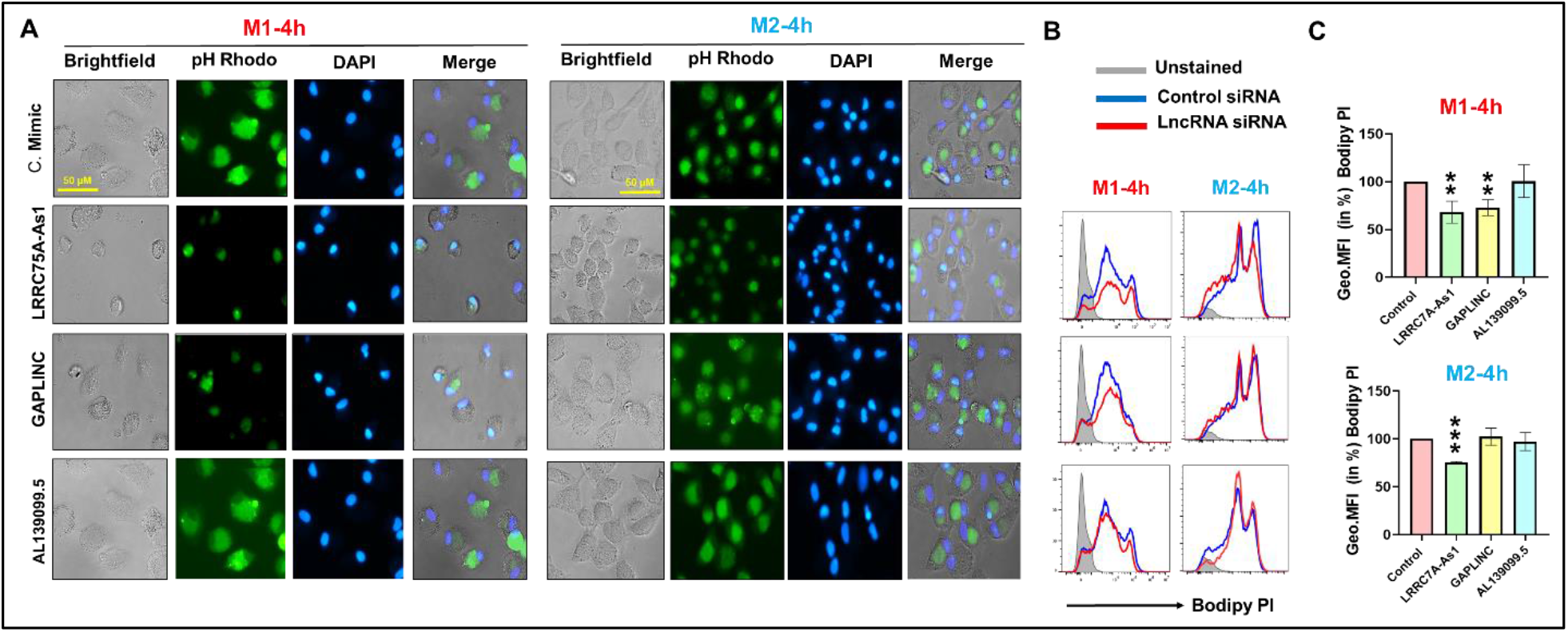
Knockdown of lncRNAs LRRC75A-As1, and GAPLINC impairs antigen processing by macrophages. **A**) M1 and M2 macrophages were transfected with control or LRRC75A-As1, GAPLINC and AL139099.5 targeting siRNA. After 36 h, cells were incubated with DQ-OVA for 4h. Representative florescent images show inhibitory impact of LRRC75A-As1, and GAPLINC on antigen processing. Final magnification: X40, scale bar corresponds to 100 μm. **B)** M1 and M2 macrophages were harvested after 4 h of DQ-OVA incubation and analyzed by flow cytometry. Overlay histograms showing differences in BODIPY dye signals in control or lncRNA targeting siRNA transfected M1 and M2 macrophages. **C)** Histograms showing percent geometrical MFI values in lncRNA-transfected M1 and M2 macrophages. The data presented as ±SEM of three independent experiments in each cell type. Student’s *t*-test was conducted to calculate *p* values. \*\**p* < 0.01, ***P < 0.001.

### Innate cytokine response in macrophages is impaired by lncRNA knockdown

Phagocytosis involves antigen recognition and activation of inflammatory responses (34–36). Next, we investigated whether secretion of the pro-inflammatory cytokines is concurrently impaired by lncRNAs. Because only LRRC75S-AS1 and GAPLINC knockdown showed biological effects on macrophage polarization, phagocytosis and antigen processing, we excluded AL139099.5 from the subsequent experiments. LncRNA knockdown M1Mφ were challenged with rhodamine labeled *E. coli* bioparticles and the supernatants were collected at 4 and 24h (**Fig. 7**). A multiplex analysis of pro-inflammatory cytokines (IL-1β, IL-6, IL-8 and TNF-α) was performed to quantify cytokine secretion profiles. Our results show that compared to control siRNA, lncRNA knockdown cells exhibit impaired pro-inflammatory cytokine levels. Overall, we reported downregulation of IL-1β and IL-6 in LRRC75S-A1 and GAPLINC knockdown but no significant changes were observed in IL-8 or TNF-α at 4h. (**Fig. 7**). Compared to control siRNA, the secretion of IL-1β and TNF-α was significantly reduced at 24h in LRRC75S-A1 and GAPLINC knockdown cells. Levels of IL-6 were downregulated in LRRC75S-AS1 RNAi cells but no significant changes were observed in GAPLINC RNAi at 24h. Interestingly, secreted levels of IL-8 were reduced in GAPLINC knockdown but not in LRRC75A-As1 knockdown at later time point (24h). In general, the downregulated secretion of pro-inflammatory cytokines by knockdown of candidate lncRNAs corroborates with the reduced phagocytosis activity.

**Fig. 7.**
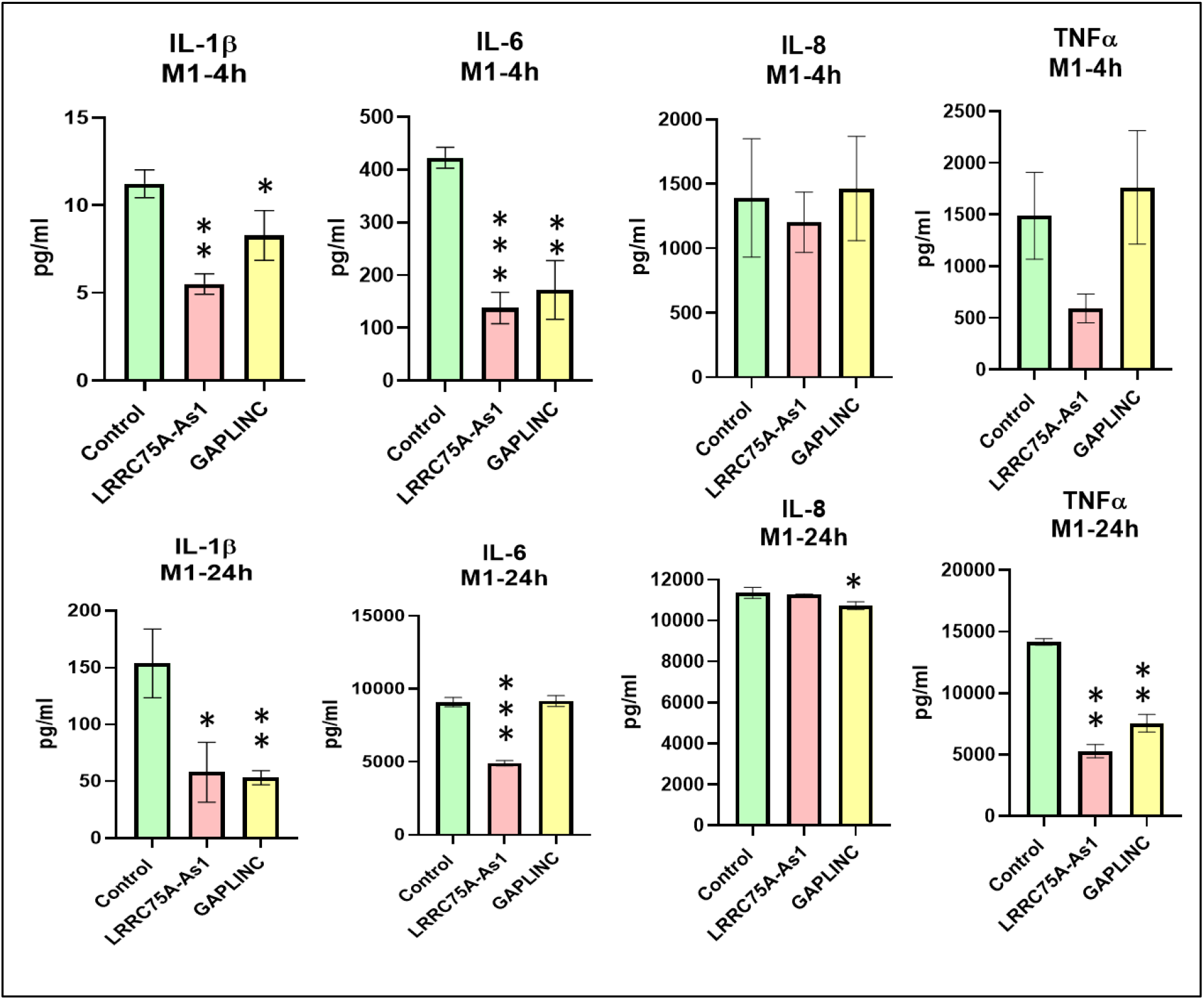
Innate immune cytokine response is impaired by lncRNA knockdown in macrophages challenged with *E. coli*. M1 macrophages were transfected with lncRNA targeting siRNA or control. After 36 h, cells were challenged with *E. coli* and the supernatants were collected at 4 and 24h. Secreted levels of IL-1β, IL-6, IL-8 and TNF-α were analyzed by multiplex bead array. Data are presented as mean ± SEM from four independent donors. Student’s *t*-test was conducted to calculate *p*-values (\**p* < 0.05, \*\**p* < 0.01, \*\*\**p* < 0.001).

### Macrophage migration in wound-healing assays is regulated by lncRNAs

Macrophage migration is essential in phagocytosing the invading pathogens/antigens, tissue patrolling, and facilitate tissue remodeling (*36–38*). These processes allow crosstalk of innate and adaptive immune cells and requires polarization of the cell body (*39*). Myeloid cell migration from tissues to lymph nodes as well as antigenic peptides generated after phagocytosis subsequently activates naïve T cells (*39, 40*). We therefore tested the impact of candidate lncRNAs on cell migration in M2Mφ. Cells were transfected with siLRRC75S-AS1 and siGAPLINC and after 36 h post-transfection we performed two functional assays to assess their impact on cell migration. In the first assay, migration was assayed by generating scratch and monitoring its healing over 48 h by capturing images of wounded area. Compared to control mimic, we observed significantly attenuated migration in LRRC75S-AS1 and GAPLINC-transfected M2Mφ. Approximately, only ∼30% of wounded area was covered after 48h of migration in siLRRC75S-AS1 and ∼27% in siGAPLINC M2Mφ (**Fig. 8A,C**). Knockdown of both the lncRNAs robustly impaired the migratory capacity of macrophages compared to control siRNA. To confirm these findings, we next performed the radius gel migration assays in cells transfected with LRRC75S-AS1, GAPLINC or control siRNA. Migration was initiated by dissolving the gel 24h post-siRNA transfection and cell migration was monitored. After 48h, wound area in control siRNA transfected cells was completely confluent while, LRRC75S-AS1 and GAPLINC knockdown M2Mφ showed a significant delay in wound closure. To quantify the migration rate, the area of the migration zone was calculated for lncRNA or control siRNA for each time point and normalized with control siRNA at the 0h time point. After 48h of migration monitoring course, LRRC75S-AS1 and GAPLINC knockdown cells showed ∼70% and ∼60% of wound remaining, which is statistically significant compared to the control siRNA (0% wound remaining) (**Fig. 8 B,D**). Impaired migration of lncRNA knockdown macrophages strongly support our previous observations of attenuated phagocytosis, antigen processing and cytokine secretion that requires efficient cell migration to perform these biological functions.

**Fig. 8.**
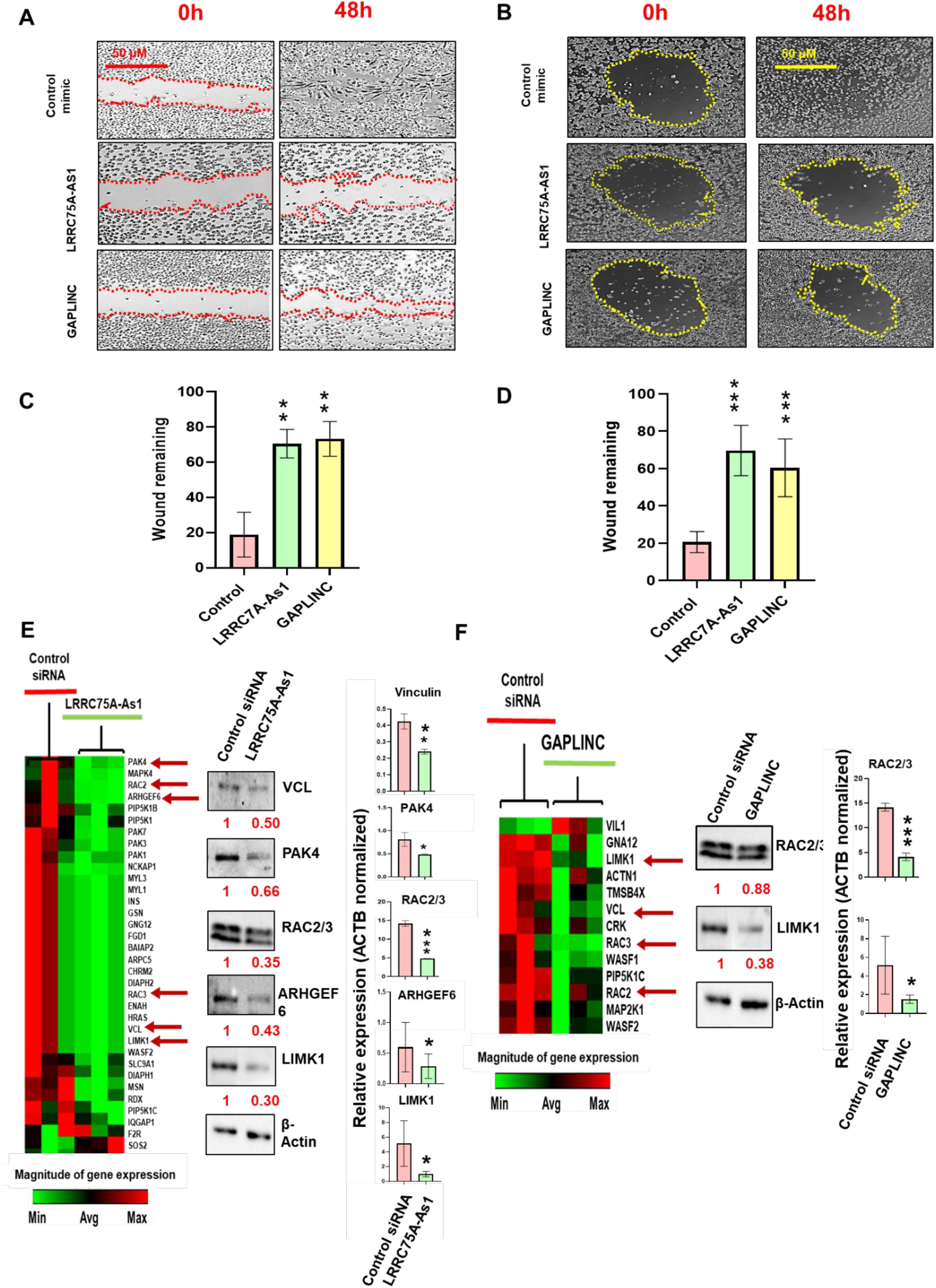
LLRC75A-As1 and GAPLINC regulate macrophage migration by modulating actin cytoskeleton signaling. **A**) M2 macrophages were transfected with control, si-LLRC75A-As1, or si-GAPLINC. After 24h of transfection, a scratch was created and cells were monitored for 48h by light microscopy. Representative images of scratched areas were performed. The red lines indicate the invasive front in the wound healing assay. Wound closure was photographed at 0h and 48 h after wounding. **B**) Macrophage migration was assessed to analyze the impact of LLRC75A-As1 and GAPLINC using wound-closure assay. Cells were transfected with siRNA targeting candidate lncRNAs and after 24 h the gel coated at the center of the well was dissolved to allow cell migration. Representative images were taken to monitor the wound closure until 48h. ImageJ analysis was performed to quantify the wound closure for 48 h in **C**) scratch and **D**) gel dissolution assays. The scratched area at control (0 h) was arbitrarily assigned as 100%. Final magnification: X100, scale bar corresponds to 100 microns. Macrophages were transfected with siRNA targeting lncRNAs and the expression of cytoskeletal pathway related genes was assessed by PCR array. Heatmaps showing differentially expressed genes in **E**) LRRC75A-As1 and **F)** GAPLINC knockdown macrophages. To validate the RT-qPCR results, protein levels of selected cytoskeletal genes (marked by red arrows in the heatmap) were quantified by Western blotting in **E**) LRRC75A-As1 (right panel) and **F)** GAPLINC (right panel) knockdown cells. Data are expressed as mean ± SEM of three independent transfections. Student’s *t*-test was conducted to calculate *p*-values (\**p* < 0.05, \*\**p* < 0.01, \*\*\**p* < 0.001).

### LRRC75S-AS1 and GAPLINC knockdown dysregulate expression of genes involved in cytoskeleton signaling

Next, we examined the underlying mechanism of attenuated migration in lncRNA knockdown macrophages. Mφ were transfected with lncRNA targeting or control siRNA and the expression of eighty-eight genes involved in actin cytoskeleton signaling was examined by PCR array. Compared to control, we identified several differentially expressed mRNAs (cut-off Ct≤35, fold change ≥1.2 or ≤0.5 and a P value ˂0.1) in lncRNA knockdown cells (**Data S4-S5**). Interestingly, LRRC75S-AS1 knockdown M2Mφ showed a higher number of differentially expressed genes compared to GAPLINC. Indeed, we observed three upregulated genes (F2R, SOS2 and SOS1) and thirty-two downregulated genes (PAK4, MAPK4, RAC2, ARHGEF6, PIP5K1B, PIP5K1, PAK7, PAK3, PAK1, NCKAP1, MYL3, MYL1, INS, GSN, GNG12, FGD1, BAIAP2, ARPC5, CHRM2, DIAPH2, RAC3, ENAH, HRAS, VCL, LIMK1, WASF2, SLC9A1, DIAPH1, MSN, RDX, PIP5K1C, IQGAP1) in si-LRRC75S-AS1 compared to control siRNA (**Fig. 8E**). In GAPLINC knockdown cells, we identified one upregulated gene (VIL1; fold change <u>˂</u> 9) and twelve downregulated genes (fold change ≥ 0.02) (GNA12, LIMK1, ACTN1, TMSB4X, VCL, CRK, RAC3, WASF1, PIP5K1C, RAC2, MAP2K1 and WASF2) (**Fig. 8F**). These results show a unique impact of each lncRNA on the genes involved in cytoskeleton signaling, with LRRC75S-AS1 being a potent regulator of the pathway.

To validate the PCR array results, we analyzed the protein expression of Vinculin (VCN), PAK4, RAC2, ARHGEF6, and LIMK1 by western blot (**Fig. 8E**). Lysates were prepared from lncRNA knockdown macrophages undergoing wound healing to analyze the expression of cytoskeleton proteins. LRRC75S-AS1 knockdown cells showed a significant reduction in the expression of Vinculin, PAK4 and RAC2/3, ARHGEF6 and LIMK1 compared to control siRNA. GAPLINC RNAi significantly reduced RAC1/2/3 and LIMK1 expression compared to control (**Fig. 8F**). These results further support a key role of LRRC75S-AS1 and GAPLINC during the migration process of macrophages to facilitate innate immune responses and prime adaptive arm of immunity.

## Discussion

Being the gate keeper of immune activities, macrophage differentiation, polarization, and functions demand a tight regulation. Dysregulated macrophage activity is implicated in a multitude of diseases *viz*. infection (*20*), fibrosis (*21*), cancer (*22*), autoimmune disease (23), insulin resistance (*24*), atherosclerosis (*25*), etc. Despite far-reaching research in the field of macrophage biology, we are still searching for the key endogenous molecules that contribute to dynamic changes in tissue microenvironment and orchestrate macrophage function towards immune homeostasis. LncRNA expression profiles in macrophage polarization is reported by multiple studies; however, their functional characterization remain less explored (41–43). Moreover, kinetics of macrophage lncRNA profiles has never been studied. This is important as lncRNA are expressed temporally and time dependent expression may allow identification of novel lncRNA sequences. in this study we have examined time-kinetics of lncRNA profiles during M1 and M2 differentiation to identify thousands of previously known and novel lncRNAs and characterized functional regulatory role of previously unannotated lncRNAs in polarization, innate immune functions viz., phagocytosis, antigen presentation, cytokine response and cell migration.

To address whether unique lncRNAs repertoire drive macrophage polarization, we differentiated monocyte to M1 and M2 macrophages and time kinetics of lncRNA expression was performed at 18h, 3d, 5d and 7d using RNA-seq. Our results show significant changes in thousands of lncRNAs during the differentiation process, and identified lncRNAs that were common and unique to M1 and M2 subsets. We noticed a remarkable changes in lncRNAs expression profiles as early as 18 h, while relatively low percentage of lncRNAs were maintained in the later stage of differentiation. These results suggests that global changes in lncRNA expression precedes acquisition of Mφ differentiation markers (commences around day 3) supporting their key role as initial responders to microenvironment changes, while the expression of select lncRNAs reflect their necessary presence in the maintenance of macrophage phenotype (M1/M2). To our knowledge, this is the first study to comprehensively perform time-kinetics of global lncRNA profiling during primary M1 and M2 macrophage differentiation. Earlier studies demonstrated lncRNA changes in monocyte differentiated macrophages (but did not focus on differentiating macrophages), and their plausible association with multiple diseases. For instance, Chen et al. studied lncRNA expressional dynamics during PMA-induced monocyte-to-macrophage and monocyte-to-granulocyte differentiation from THP-1 and HL-60 cells (human promyelocytic leukemia cell line), respectively. They noticed differential changed in 390 lncRNAs and identified lnc-MC as a highly abundant lncRNA in the differentiated macrophages but not in granulocytes (*44*). In other study, Yang et al. characterized the role of lncRNA *NTT* in resting human primary monocyte, monocyte-derived macrophage and the THP-1 cell line. Studying disease severity and untreated patients of Rheumatoid Arthritis (RA), they reported ∼10 fold increase in the expression of lncRNA NTT in macrophages obtained from untreated RA patients (*45*). Furthermore, in a different study, Müller and colleagues showed that lncRNAs derived from CD14+ monocytes from RA patients respond to antibody treatment. They showed an upregulation of 85 lncRNAs after treating RA patients with anti-tumour necrosis factor (TNF)-α (adalimumab) or anti-interleukin (IL)-6R (tocilizumab) therapy (*46*) implying a role of lnRNAs in monitoring macrophage activity.

While a large number of lncRNAs have been identified in macrophage differentiation and the list continue to grow; majority of them remain functionally uncharacterized. In this study, we focused on three upregulated lncRNAs LRRC75A-AS1, GAPLINC and AL139099.5 that exhibit similar expression pattern in both M1/M2 subsets, albeit with varied degree of fold changes. Differential upregulation of these lncRNAs from 18h to 7d strongly suggest their function as initial responder to *in vitro* differentiation cues, which may be required to facilitate the polarization process necessary to maintain the immune homeostasis. Interestingly, LRRC75A-AS1, GAPLINC and AL139099.5 were uniquely enriched in M1 and M2 macrophages but exhibit low or undetectable expression in monocyte-derived DCs indicating their functional role in macrophage biology. Earlier, we have reported the surface expression of CDw93 and CD68 is the reflection of monocyte-to-macrophage differentiation (*47*). Keeping this in view, we have explored the role of candidate lncRNA in macrophage differentiation. RNAi studies revealed that knockdown of LRRC75A-AS1 and GAPLINC, but not AL139099.5, significantly downregulated surface expression of macrophage differentiation markers CDw93 and CD68. Studies demonstrating the role of lncRNA in macrophage differentiation are scarce. Yang et al. demonstrated that lncRNA NTT regulate CD68 expression during monocyte-to-macrophage differentiation process (*45*). Reports from our lab and others also described the role of RN7SK, GAS5 and MALAT1 in macrophage differentiation (*42,43*). Studies from our lab reported induction of RN7SK and GAS5 in monocyte-to-M2Mφ differentiation. Knockdown of RN7SK and GAS5 exhibit downregulation of M2 surface markers (CD163, CD206 or Dectin) and concomitant induction of M1 markers (MHC II or CD23) indicating pro-M2 functions. This data clearly suggests that lncRNAs are responsive to the extracellular milieu and might be involved in regulating macrophage polarization (*42*). Furthermore, Cui et al., reported the expressional increase of lncRNA MALAT1 in both human PBMCs derived and mouse bone marrow–derived macrophages (BMDMs). Interestingly, they reported that MALAT1 expression is needed to control early events (4h to 24h) and late events of macrophages differentiation in mouse and human macrophages respectively (*48*). Consistent with this, Ahmad et al. reported MALAT1 upregulation in both M1 and M2 Mφ. Knockdown of MALAT1 upregulated M2 markers without affecting M1 marker levels demonstrated its function in skewing Mφ polarization (*43*). Together, these studies show the relevance of lncRNA expression towards the monocyte-to-macrophage differentiation; however, LRRC75A-AS1, and GAPLINC characterized here appears to be more relevant because these lncRNAs are needed throughout the differentiation process and exhibit macrophage-specific expression, reflecting their essential role in the process.

After differentiation, macrophages need to be polarized to inflammatory (M1) phenotype (to eliminate the pathogen) or perform reparative (M2) function (towards the disease resolution). Knockdown of LRRC75A-AS1 and GAPLINC reduced M1 macrophage polarization markers HLA-DR and CD32. Thus, the expression of these lncRNAs during monocyte to M1 and M2 differentiation seemingly regulate the macrophage polarization events. So far, only few studies suggest the role of lncRNAs in macrophage polarization. Huang et al. elucidated that overexpression of LncRNA TCONS_00019715 in M1Mφ induced by IFNγ and LPS) rather than IL-4 induced M2 Mφ (*11*). Another study demonstrated that lncRNA FENDRR plays imperative role in the polarization of IFNγ-stimulated M1 macrophages (*49*). Furthermore, during the pathogenesis of experimental autoimmune encephalomyelitis (EAE), the upregulation of lncRNA GAS5 was associated with impaired M2 polarization (*50*). Recently published study from our lab also deciphered the association between lncRNAs RN7SK and GAS5 expression and M1 and M2 polarization (43). Interestingly, studies also suggested the association between lncRNA expression dynamics during TLR activation and resultant NF-kB signaling, which are related to M1 polarization of Mφ in inflammatory microenvironment. LncRNAs Rel (*51*) and FIRRE (*52*) are upregulated during LPS-induced TLR activation of macrophages, which might eventually result towards M1 polarization after epigenetically activating the immune response protein genes located in proximity to these lncRNAs. Interestingly, simultaneous upregulation lncRNAs that promotes as well as negatively regulates NF-κB signaling upon TLR-activation: lncRNA-Cox2, lncRNA-AK170409 and lncRNA-Cox2, respectively in the macrophages, suggest that lncRNAs possess capability of regulating immune homeostasis via controlling TLR-dependent downstream signaling in macrophages that could otherwise result in excessive immune mediated tissue damage (*53*).

Antigen uptake/processing, phagocytosis and immunosurveillance are key functional attributes of macrophages (*35,36*). LncRNA regulation of antigen uptake/phagocytosis in macrophages is an understudied topic and only few reports demonstrate lncRNA regulation of these central macrophage functions. Das et al., demonstrated that overexpression of lncRNA *Dnm3os* in facilitates inflammation and immune response to promote the phagocytosis in macrophages (*54*). In another study, Hung et al, showed that the knocking down of plaque-enriched lncRNA in atherosclerotic and inflammatory bowel macrophage regulation (PELATON) lncRNA; evidently reduced the phagocytosis in monocyte-derived macrophages by reducing the CD36 mRNA (*55*). Previously, we have shown that siRNA mediated knockdown of GAS5 and RN7SK increase antigen uptake and processing in both M1 and M2 Mφ (*42*). In this study, we demonstrated attenuated bacterial phagocytosis and ovalbumin uptake as well as processing in M1 macrophages treated with siLRRC75A-AS1 reflecting their differential regulation of macrophage biology. Compared with monocytes, macrophages exhibit higher antigen uptake/phagocytosis capacity. Since our candidate lncRNAs hampers the differentiation of monocyte-to-M1/M2 Mφ, therefore, impaired phagocytosis and antigen uptake is expected upon LRRC75A-AS1 and GAPLINC knockdown. Therefore, the present study in conjunction with above reports strongly suggested the regulation of both phagocytosis and antigen uptake of macrophages via lncRNAs.

Cytokine secretion is a key macrophage activity towards pathogen clearance and disease resolution. M1 polarized macrophages secrete various pro-inflammatory cytokines that play key role in conflagration of T cells for the activation of adaptive immune system (*32–34*). In our study, we observed significantly higher secretion of IL-1β, IL-6, IL-8 and TNFα in macrophages, which was dampened in LRRC75A-AS1 and GAPLINC knockdown strongly supporting the role of lncRNAs in cytokine secretion and in line with our results showing attenuated bacterial phagocytosis. Multiple studies have reported lncRNA regulation of pro-inflammatory cytokines/gene expression. For instance, Sun et al. demonstrated that lncRNA GAS5 overexpression favors M1 phenotype by inducing nitric oxide synthase (iNOS), IL-1β, and TNF-α. Similarly, GAS5 augments proinflammatory cytokines (IL-6, IL-1β, and TNF-α) levels in ox-LDL-induced THP-1 macrophages (50). Interestingly, lncRNA HOTAIR regulates the functions of both M1 and M2 Mφ. The treatment of primary macrophages and BMDMs with LPS induces the HOTAIR, which mechanistically favors the degradation of IκBα and promote nuclear translocation of NF-κB to activate the pro-inflammatory cytokine genes. On the contrary, in the tumor microenvironment HOTAIR facilitates the expression of CCL2 in this lncRNA in macrophages/myeloid-derived suppressor cells (MDSCs), which eventually induces tumor growth and metastasis (*56*). Munteanu et al. showed that stimulation of THP1 cells by IFN-γ, induces the FENDRR and M1 macrophage-specific cytokines including IL-1β, TNF-α, and CXCL10. Enhanced phosphorylation of STAT1 is considered as one of plausible mechanism of pro-inflammatory cytokine induction (*49*). LncRNA MALAT1 induction was reported by LPS stimulation in PMA-induced THP-1 and RAW264.7 cells (*57*). Interestingly, MALAT1 knockout ameliorates the LPS-induced activation of M1 macrophages, by negatively regulating NF-κB signaling pathway supporting regulatory function of MALAT1 in inflammatory cytokines expression. However, the mechanism through which lncRNAs LRRC75A-AS1 and GAPLINC regulate the cytokine signaling needs further investigation.

Macrophages constantly patrol tissues for antigens and represent one of the earliest innate immune cells to infiltrate infected sites the influence of local inflammatory mediators by virtue of their immense potential to rapidly and transiently modify their cellular architecture (*58–60*). Therefore, cytoskeletal homeostasis associated pathways must play crucial roles in regulating these migratory events (*7*) and therefore need to be tightly regulated. Studies from our lab and others strongly suggest the impairment in the actin rearrangement or organization is responsible for attenuated macrophage migration (*61–63*). Scientific reports demonstrating the regulatory roles of lncRNAs associated with cell migration are mainly limited to cancer invasion and metastasis. LncRNA PCAT-1 upregulation in prostate tissues and cells lines is associated with increased migration, proliferation, and invasion of PCa cells (*64*), while induction of lncRNA ZFAS1 plays a critical role in increasing the proliferation, invasion, and migration of epithelial ovarian cancer cell lines (*65*). Furthermore, high expression of HOTAIR is associated with advance pathological stage, poor tumor differentiation, increased tumor progression, metastasis, and poor OS (*66,67*). The role of lncRNA expression on the migration of macrophages remain understudied. In this study, we show that LRRC75S-AS1 and GAPLINC knockdown Mφ exhibit impaired migration as observed by wound healing assays. To the best our knowledge, this is first evidence in scientific literature showing the association of lncRNA with the macrophage migration. Mechanistically, knockdown of these lncRNAs showed the reduced expression of key proteins (Vinculin, PAK4 and RAC2/3, ARHGEF6 and LIMK1) involved in macrophage migration. Our results show mechanisms underlying novel functions of LRRC75S-AS1 and GAPLINC in regulating cytoskeletal genes that are essential for macrophage migration critical in performing innate immune functions including phagocytosis, antigen uptake, cytokine secretion, etc. In conclusion, we have performed the time-kinetics of global lncRNA profiles during monocyte-to-macrophage (M1/M2) differentiation (18h, 3d, 5d, 7d) and identified thousands of previously reported and novel lncRNAs, and demonstrated functional role of lncRNAs, expressed selectively in macrophages, in cell differentiation and polarization. We have characterized functionally unannotated lncRNAs LRRC75S-AS1 and GAPLINC in regulating key innate immune functions including antigen uptake, phagocytosis and cytokine secretion. Mechanistically, these lncRNAs modulate various genes that regulate cytoskeletal dynamics linking their role in functional immune activity. Overall, our data provide a rich resource to study macrophage-specific lncRNAs and characterized novel functional roles of LRRC75S-AS1 and GAPLINC in macrophage biology.

## Materials and Methods

### Primary human monocyte isolation and differentiation

Freshly prepared buffy coats were collected from healthy donors (Sylvan N. Goldman, Oklahoma Blood Institute, Oklahoma City, OK, USA) and CD14+ monocytes were obtained by density gradient centrifugation and magnetic bead isolation. In brief, PBMCs were purified by use of Ficoll Paque (GE Healthcare, Piscataway, NJ, USA)-based density centrifugation. PBMCs were incubated with magnetically labeled CD14 beads (Miltenyi Biotec, Cologne, Germany), according to the manufacturer’s instructions. Monocyte purity and viability were 95%, as determined by flow cytometry. Monocytes were plated at a density of 2X10^6^/ml in DMEM, supplemented with penicillin (100 U/ml), streptomycin (100 mg/ml), and gentamicin (50 mg/ml). After 2 h, the media were substituted with media containing 10% heat-inactivated FBS (Life Technologies, Grand Island, NY, USA) and rhGM-CSF or rhM-CSF (both 50 ng/ml; PeproTech, Rocky Hill, NJ, USA) for M1 and M2 macrophage, respectively. Media were replaced every 72 h. Cells were harvested at 18h, day 3, 5, and 7 for total RNA isolation. At day 7, cells were harvested and differentiation confirmed by flow cytometric analysis of CDw93, CD68, CD11b, and CD14 expression.

### Library construction and sequencing

Total RNA was extracted using miRNeasy kit (Qiagen, CA, USA) following the manufacturer’s procedure. The total RNA quality and quantity were analyzed by Bioanalyzer 2100 and RNA 6000 Nano LabChip Kit (Agilent, CA, USA) with RIN number >7.0. Approximately 1 ug of total RNA was used to remove ribosomal RNA according to the manuscript of the Epicentre Ribo-Zero Gold Kit (Illumina, San Diego, USA). Following purification, the ribo-minus RNA fractions is fragmented into small pieces using divalent cations under elevated temperature. Then, the cleaved RNA fragments were reverse-transcribed to create the final cDNA library in accordance with a strand-specific library preparation by dUTP method. The average insert size for the paired-end libraries was 300±50 bp. Next, pair-end 2×150bp sequencing was performed on an illumina Hiseq 4000 platform housed in the LC Sciences (Hangzhou, China) following the vendor’s recommended protocol.

## Bioinformatics analysis

### Transcripts Assembly

Firstly, Cutadapt (*68*) and perl scripts in house were used to remove the reads that contained adaptor contamination, low quality bases and undetermined bases. Then sequence quality was verified using FastQC (http://www.bioinformatics.babraham.ac.uk/projects/fastqc/). We used Bowtie2 (*69*) and Tophat2 (*70*) to map reads to the genome of Homo sapiens (Version:). The mapped reads of each sample were assembled using StringTie (*71*). Then, all transcriptomes from 22 samples were merged to reconstruct a comprehensive transcriptome using perl scripts and gffcompare (https://github.com/gpertea/gffcompare/). After the final transcriptome was generated, StringTie (*71*) and Ballgown (*72*) was used to estimate the expression levels of all transcripts.

#### LncRNA identification

For lncRNAs identification, transcripts that overlapped with known mRNAs, known lncRNAs and transcripts shorter than 200 bp were discarded. Then we utilized CPC (*28*) and CNCI (*29*) to predict transcripts with coding potential. All transcripts with CPC score <-1 and CNCI score <0 were removed. The remaining transcripts with class code (I, j, o, u, x) were considered as lncRNAs.

#### Different expression analysis to transcripts

StringTie (*71*) was used to perform expression level for mRNAs and LncRNAs by calculating FPKM (FPKM = [total_exon_fragments / mapped_reads (millions) × exon_length (kB)]). The differentially expressed mRNAs and LncRNAs were selected with log2 (fold change) >1 or log2 (fold change) <-1 and with parametric F-test comparing nested linear models (p value < 0.05) by R package Ballgown (*72*).

#### Target gene prediction and functional analysis of LncRNAs

To explore the function of lncRNA, we predicted the cis-target genes of lncRNAs. LncRNAs may play a cis role acting on neighboring target genes. In this study, coding genes in 100,000 upstream and downstream were selected by perl script. Then, we showed functional analysis of the target genes for LncRNAs by using the scripts in house. Significance was expressed as a p value < 0.05.

#### Localization of lncRNAs on genome

Genomic localization and abundance of lncRNAs was examined using the program Circos (*56*). The lncRNAs were subdivided into five categories according to their class code generated by StringTie: (i) a transfrag falling entirely within a reference intron (intronic); (j) potentially novel isoform or fragment at least one splice junction is shared with a reference transcript; (o) generic exonic overlap with a reference transcript; (u) unknown, intergenic transcript (intergenic); (x) Exonic overlap with reference on the opposite strand (antisense). Those transcripts assembled by StringTie which are not annotated in genome annotation database are belong to novel transcripts with class code (i, j, o, u, and x).

### Total RNA extraction, cDNA Isolation and qPCR analysis

Cells were washed three times with PBS, and 700 µl of TriZol reagent (Invitrogen, CA, USA) was added to 24-wells culture plate in each condition. Total RNA was isolated from 18 h, day 3, day 5, and day 7 differentiated cells using miRNeasy micro kit (Qiagen, Gaithersburg, MD, US). A total of 250 ng RNA was used to synthesize cDNA, which was synthesized from high capacity cDNA Reverse transcription kit (ThermoFisher Scientific,

Grand Island, NY, USA). The expression levels of LRRC75A-AS1, GAPLINC, LINC01010, and beta-actin genes were analyzed by qPCR reaction using SYBR Green Gene Expression Master Mix (Applied Biosystems, USA) in a StepOne 7500 thermocycler (Applied Biosystems, USA). The Ct values of three replicates were analyzed to calculate fold change using the 2^−ΔΔCt^ method.

### Transient siRNA Transfection

Monocytes were plated at 24 well/plate and treated with rhGM-CSF or rhM-CSF during 7 days (both 50 ng/ml; PeproTech, Rocky Hill, NJ, USA) for M1 and M2 macrophage differentiation, respectively (as described above). Transient transfection of siRNA was performed in MΦM1 and MΦM2, respectively, using Lipofectamine 2000 (Invitrogen-Life Technologies Corporation, Carlsbad, CA, USA), as per manufacturer’s instructions. LncRNAs targeting siRNAs targeting LRRC75A-AS1, GAPLINC, LINC01010 transcripts were designed by IDT siRNA design tool and control siRNA, were purchased from Integrated DNA Technologies Inc. (Coralville, Iowa). After 36 h of transfection, total RNA was using miRNeasy micro kit (Qiagen, Gaithersburg, MD, US). A total of 250 ng RNA was used to synthesize cDNA, which was synthesized from high capacity cDNA Reverse transcription kit (ThermoFisher Scientific, Grand Island, NY, USA). The expression levels of LRRC75A-AS1, GAPLINC, LINC01010, and beta-actin genes were analyzed by qPCR reaction using SYBR Green Gene Expression Master Mix (Applied Biosystems, USA) in a StepOne 7500 thermocycler (Applied Biosystems, USA). The Ct values of three replicates were analyzed to calculate fold change using the 2^−ΔΔCt^ method. siRNAs were used at a final concentration of 50 nM. As a positive control, a fluorescent oligonucleotide duplex labeled with DY-547, siGLO Red Transfection Indicator, was used (ThermoFisher Scientific, GrandIsland, NY, US). After 36 h post-transfection, M1 and M2Mφ were treated with TriZol reagent (Invitrogen, CA, USA).

### Flow Cytometry

Cells were harvested and washed in ice-cold PBS supplemented with 1% (v/v) FBS and 0.08% sodium azide. Cellular debris were excluded based on size (forward scatter [FSC]) and granularity (side scatter [SSC]). The FSC/SSC gate for Monocyte comprised ∼60%, total events. Couplets were excluded based on SSC versus FSC and SSC versus pulse width measurements. M1 and M2Mφ were stained for cell surface markers with FITC, PE, and APC conjugated antibodies. For polarization analysis, human antibodies for CD32-PE, HLA-DR-FITC (both M1 marker), CD163-APC, CD206-PE (both M2 marker) were purchased from BD Pharmingen (San Diego, CA, USA) or BioLegend (San Diego, CA, USA). Unstained and isotype control (BD Pharmingen (San Diego, CA, USA) were used as controls. Samples were analyzed using a BD Accuri™ C6 flow cytometer (BD Biosciences, San Jose, CA. Further analysis was performed using FlowJo software (Tree Star, Ashland, OR). Cells were gated according to their forward scatter (FSC) and side scatter (SSC) properties including the larger cells with high granularity and excluding the small-sized debris with a low SSC and FSC shown at the bottom left corner of the dot plot.

### Phagocytosis Assay

Monocytes-derived M1 and M2 macrophages (400,000/well, 96-well plate) were transfected on day 7 with LncRNAs -siRNAs (LRRC75A-AS1, GAPLINC and Al139099.5) or control siRNA. Phagocytosis assay was performed with pHRodo Green conjugated E. coli (Invitrogen, Carlsbad, CA) 24 h post-transfection, according to the manufacturer’s protocol. Briefly, the labeled E. coli bioparticles were resuspended in Live Imaging Buffer (Life Technologies) at final concentration of 1 mg/ml and homogenized by sonication for 2 min and resuspended in culture media. Cells were incubated with labeled *E. coli* for 2 h at 37 °C, then washed three times with PBS and fixed with 4% paraformaldehyde. Finally, images were captured from four random fields per well per donor using EVOS florescent microscope (ThermoFisher Scientific, GrandIsland, NY, US) and the Rhodamine expression was analyzed by flow cytometry.

### Cytokine Analysis

M1 and M2Mφ transfected with LRRC75A-AS1, GAPLINC, Al139099.5 or control siRNA were challenged with pHRodo Green conjugated *E. coli* and supernatants were collected after 4 and 24 h. The cytokine/chemokine levels were analyzed by multiplex assays.

Multiplex analysis of IL-6, IL-8, IL-1β and TNF-α was performed using Milliplex MAP Human Cytokine/Chemokine Magnetic Bead Panel (Millipore, Billerica, MA, USA). Data were collected on Bio-Plex Flow cytometer (Bio-Rad, Hercules, CA, USA).

### Antigen Uptake and Processing Assay

Monocytes-derived M1 and M2 macrophages (400,000/well, 96-well plate) were transfected with siRNA targeting LRRC75A-AS1, GAPLINC, AL139099.5 or control siRNA. After 36 h post-transfection, cells were treated with DQ^TM^-conjugated Ova (1mg/ml, Molecular Probes, Grand Island, NY) in respective complete media for 2 h at 37°C. DQ^TM^-Ova consist of Ova that are heavily conjugated with BODIPY FL, resulting in self-quenching. Upon proteolytic degradation of DQ-Ova to single dye-labeled peptides, bright green fluorescence is observed. After assay incubation, cells were washed thrice with PBS, fixed with 2% PFA and Bodipy FL expression was analyzed by flow cytometry. Also, six random field images were captured on EVOS florescent microscope (ThermoFisher Scientific, GrandIsland, NY, US) for each donor (n=3) to show the BODIPY florescence expression in siRNA-transfected cells.

### Wound healing assay

Primary Mφ were grown on 48 well/plates to a nearly confluent monolayer and differentiated into M2 phenotype for 7 days. After 24 h post-transfection with LRRC75A-AS1, GAPLINC, Al139099.5 or control siRNA, the confluent monolayers were scratched to form a “wound” using a sterile needle (20 gauge × 1.5 in.). Cellular debris were removed by washing with PBS. The images were recorded for a period of 48h to monitor the migration of cells into the wounded area using a light photomicroscope at 10X magnification. To quantify (Image-J software) the percentage of wound (scratch area) at 0 h (control) was arbitrarily assigned as 100% and the percentage of wound healing at 48h was compared to control. Each assay was performed in triplicate.

### Cell migration assay

Cell migration assay was performed using the Radius™ Cell Migration Assay Kit (Cell Biolabs, San Diego, CA, USA) following the manufacturer’s instructions. Monocytes-derived M2 macrophages were seeded in 24 well plate at a density of 100,000/well and transfected with si-LRRC75A-AS1, si-GAPLINC, si-AL139099.5 or control siRNA at 50 nM. After 36 h of transfection, the biocompatible hydrogel spot was removed and the cell-free area was exposed for cell migration. Cells were monitored for 48h and images were captured on EVOS microscope (ThermoFisher Scientific, Grand Island, NY, USA) at 4× magnification. ImageJ software was used to quantify would healing changes. The wound scratch area at 0h, for LRRC75A-AS1, GAPLINC, AL139099.5 and control siRNA was arbitrarily assigned as 100% and the percentage of wound healing was calculated compared to 0h for every condition. Each assay was performed in triplicate.

### PCR array targeting cytoskeletal rearrangement related genes

Monocytes-derived M2 macrophages transfected with siRNAs targeting LRRC75A-AS1, GAPLINC, AL139099.5 or control. After 36 h, a scratch was created using sterile needle and the cells were harvested after 24 h. Total RNA was isolated using the miRNeasy Micro Kit (Qiagen), according to manufacturer’s instructions. First-strand cDNA was synthesized from 500 ng total RNA using the High-Capacity cDNA Reverse Transcription kit (ThermoFisher Scientific, Grand Island, NY, USA). A custom PCR array plates (96 well), containing 88 primer sets directed against human actin cytoskeleton signaling related genes and 8 housekeeping primer sets (Real Time Primers, LLC Elkins Park, PA). One microgram of cDNA was aliquoted onto each well, and real-time PCR was performed using a StepOne 7500 thermocycler (Applied Biosystems, Carlsbad, CA). PCR array data were analyzed using the Qiagen GeneGlobe Data Analysis Center (https://www.qiagen.com/us/shop/genesand-pathways/data-analysis-center-overview-page/). Expression levels were normalized with respect to beta-2 microglobulin (B2M) as housekeeping as it demonstrated the most consistent levels across all transfections. Next, the fold change was calculated with respect to the control siRNA. Real-time PCR was also carried out in three independent donors.

### Immunoblotting

Monocytes-derived M2 macrophages transfected with siRNAs targeting LRRC75A-AS1, GAPLINC, AL139099.5 or scramble control. After 36 h, a scratch was created using sterile needle and the cells were harvested after 24 h. The cell pellet was incubated for 15 min on ice with 300μl of cell lysis buffer (Cell Signaling Technology Incorporated, Danvers, MA, USA) with protease and phosphatase inhibitor cocktail (ThermoFisher Scientific, Grand Island, NY, USA) and centrifuged at 10,000 ×g for15 min at 4 °C. Total protein concentration was quantified by a standard Bradford assay using the colorimetric reagent from BioRad Laboratories (Hercules, CA, USA). Proteins (25μg) were separated onto SDS polyacrylamide gels using a 4–12% Bis-Tris gradient gels in the BioRad Criterion System and transferred to nitrocellulose membranes, which were blocked with 3% BSA, and probed with the appropriate antibodies. For western blot analysis, the following antibodies were used: Vinculin antibody, Pak4 (G222) Antibody, LIMK1 Antibody, Rac 1/2/3 Antibody, Cofilin (D3F9) XP Rabbit mAb, Cool2/alpha-Pix (C23D2) Rabbit mAb; all from Cell Signaling Technology Incorporated. β-Actin Antibody (C4), mouse IgG HRP-linked antibody and rabbit IgG HRP-linked antibody were purchased from Santa Cruz Biotechnology Inc. (Dallas,TX, USA). Immunocomplexes were detected with appropriate horseradish peroxidase-conjugated secondary antibodies and detected by enhanced chemiluminescence with the ECL Plus Western Blotting Detection System or ECL Advance Western Blotting Detection Kit (GEHealthcare Life Sciences, Little Chalfont, UK).Images were captured on a ChemiDoc XRS Imaging System (BioRad).

### Statistical Analysis

All the data were analyzed and plotted using GraphPad Prism (La Jolla, USA). The results are represented as SD or SEM from three independent replicates. P-values were calculated using Students t-test, and P <0.05 was considered significant. *P < 0.05, **P < 0.01, ***P < 0.001.

## Acknowledgments

This work was funded by the NIH/NIDCR RO3 DE027147 and RO1 DE027980 to ARN.

**Author contributions: Araceli Valverde**: Writing-original draft, performing experiments and data analysis, figure preparation and editing; **Raza Ali Naqvi**: Writing-original draft, performing experiments and data analysis; **Afsar Naqvi**: Conceptualization, writing-original draft preparation, editing, and supervision.

**Competing interests:** The authors declare that they have no competing interests.

**Data and materials** availability: All data needed to evaluate the conclusions in the paper are present in the paper and/or the Supplementary Materials.

