## Supplemental Figures and Tables for "Global Profiling of Differentiating Macrophages Identifies Novel Functional Long Non-coding RNAs Regulating Polarization and Innate Immune Responses"

**SUPPLEMENTARY FIGURES**

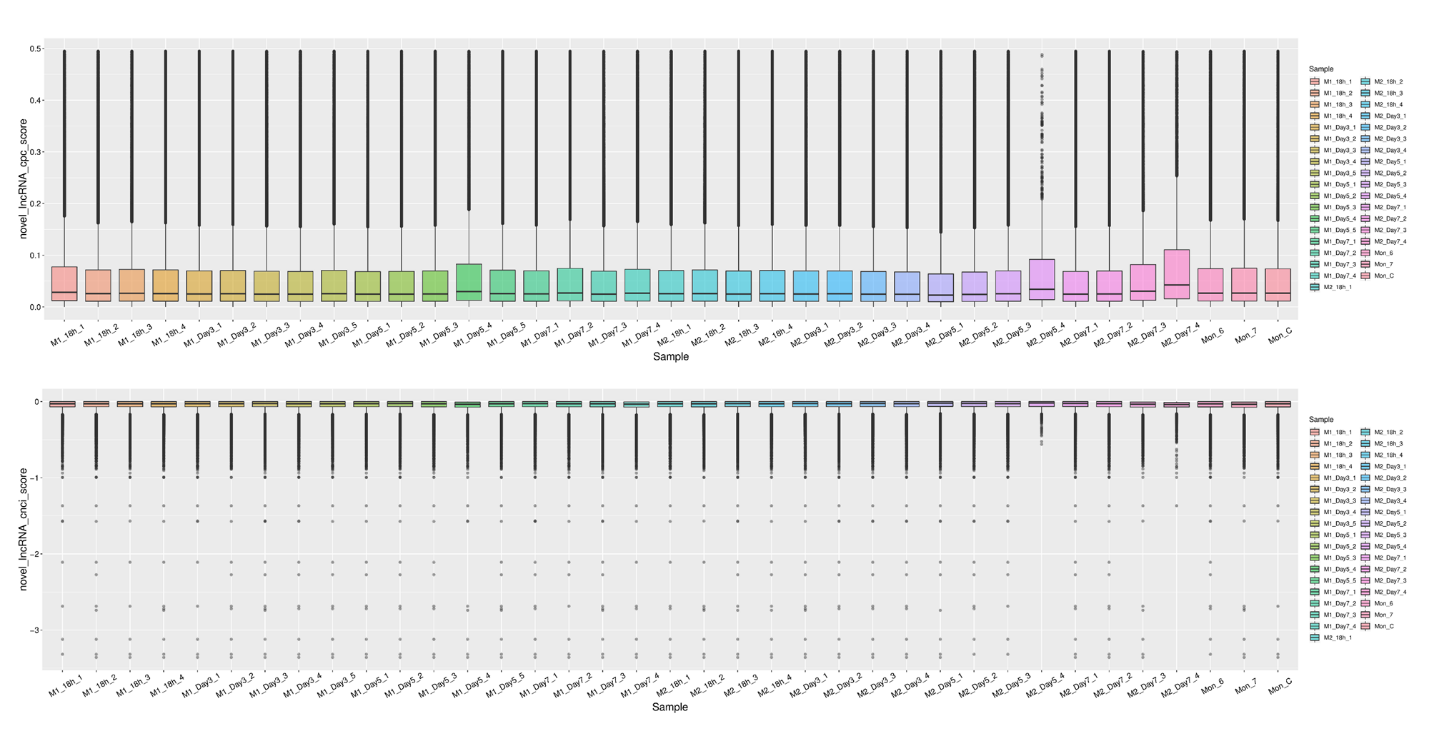

Fig. S1. Statistics of CPC and CNCI scores. Unqualified transcripts with length <200nt and coverage (count number) <3 were removed and the coding transcripts and known

lncRNA were annotated by CPC (Coding Potential Calculator) and CNCI (Coding-Non-Coding Index to predict coding potential of those transcripts.

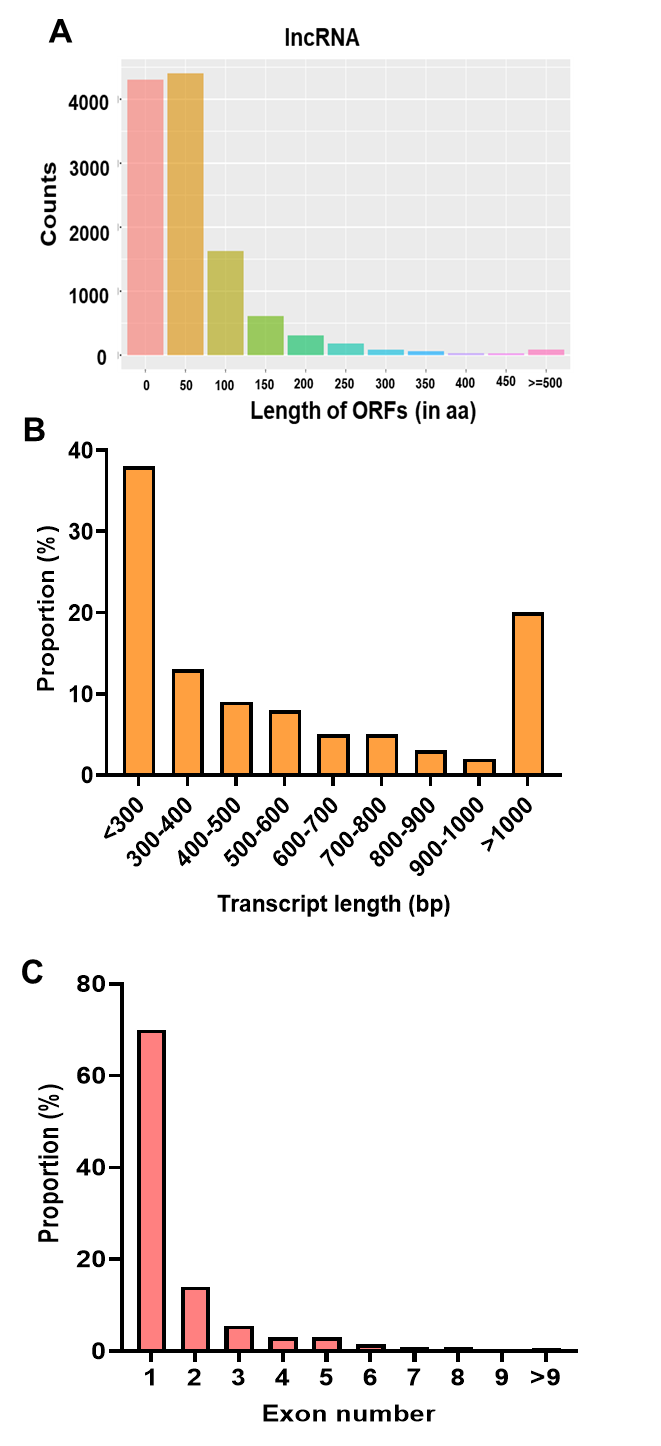

Fig. S2. Key features of lncRNA sequences identified in RNAseq. (A) ORF length, (B) Transcript length, and (C) Exon numbers of lncRNAs detected in differentiating macrophages.

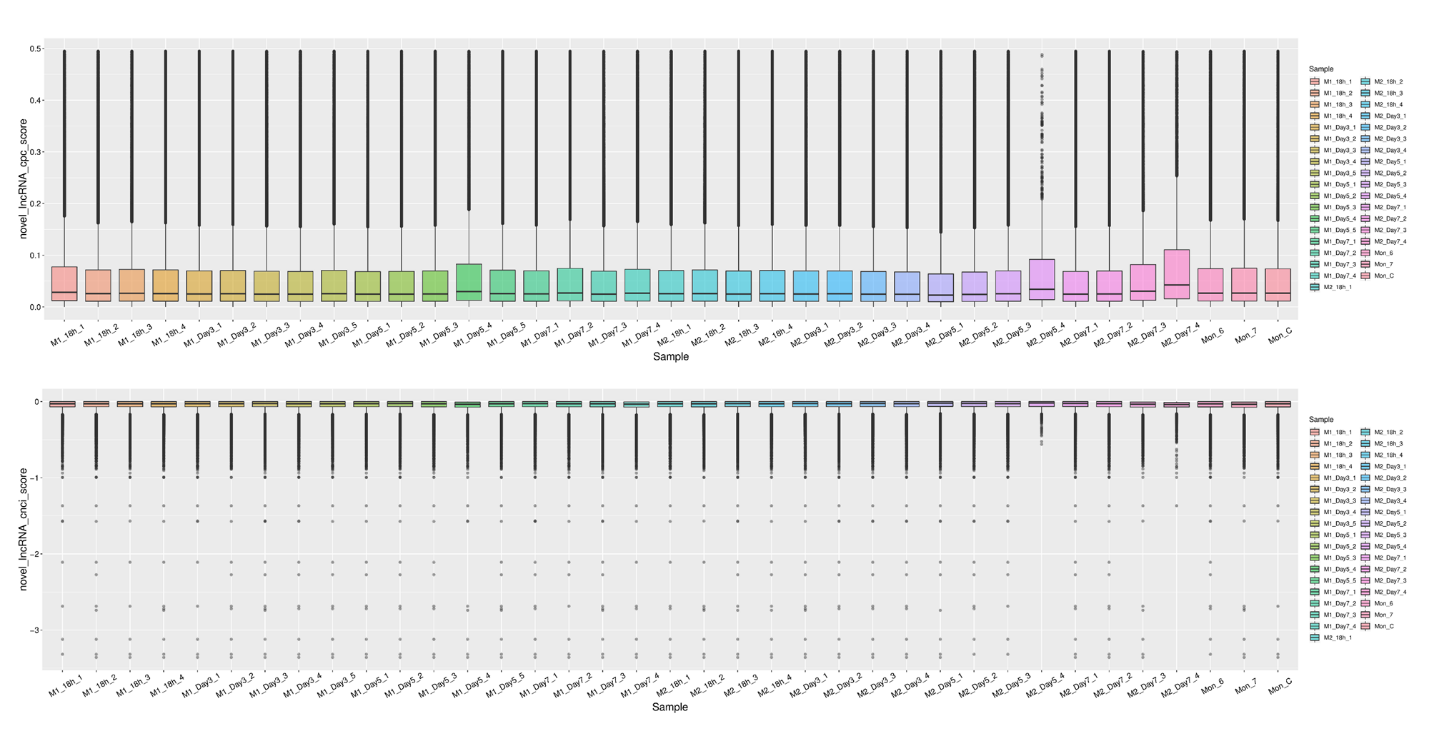
Fig. S3. Statistics of CPC and CNCI scores. Unqualified transcripts with length <200nt and coverage (count number) <3 were removed and the coding transcripts and known

lncRNA were annotated by CPC (Coding Potential Calculator) and CNCI (Coding-Non-Coding Index) to predict coding potential of those transcripts.

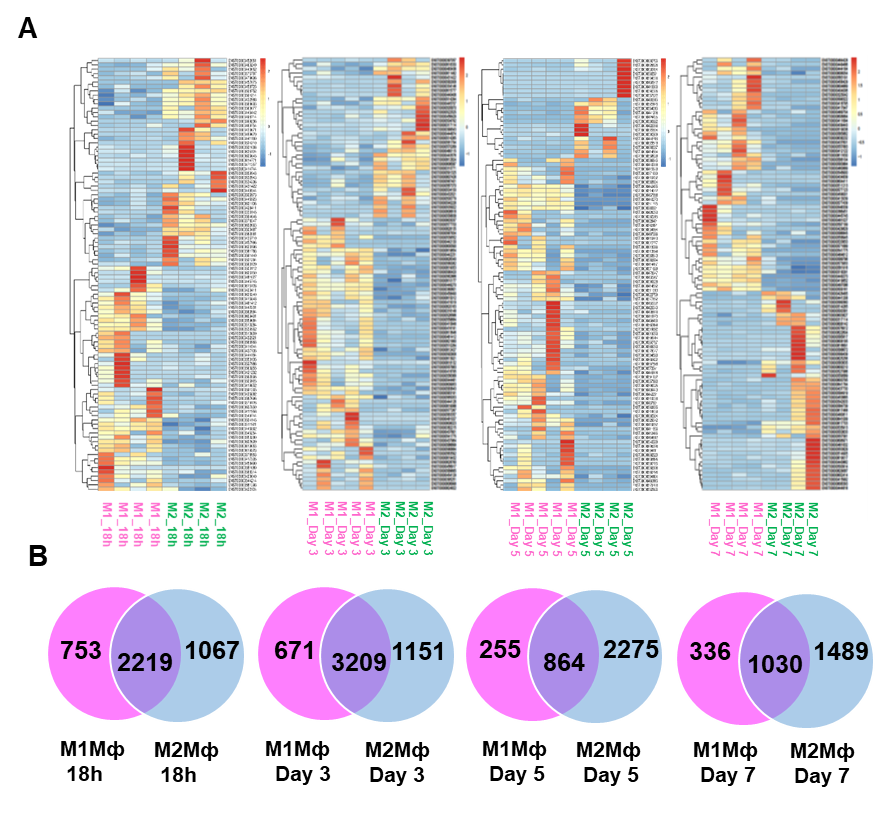

**Fig. S4**. **LncRNA differential expression profile occur during macrophage polarization. (A)** Heatmaps showing expression profiles of selected upregulated and downregulated lncRNA between M1 and M2 at 18h, day 3, 5 and 7. **(B)** Venn diagram showing the number of common and unique lncRNAs expressed in M1 and M2 at 18h, day 3, 5 and 7.

SUPPLEMENTATY TABLES

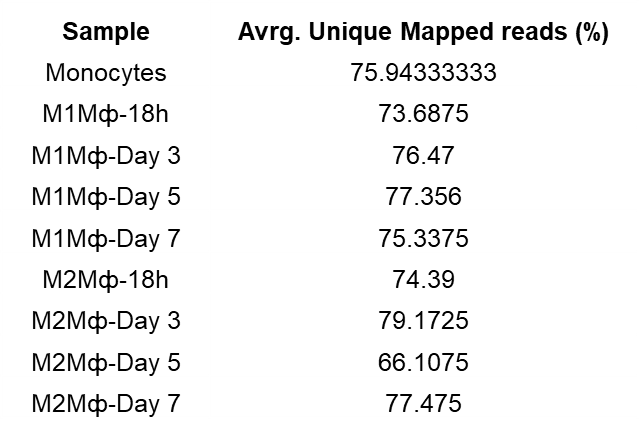

**Table S1.** Summary of average of unique mapped reads.

SUPPLEMENTARY FILES

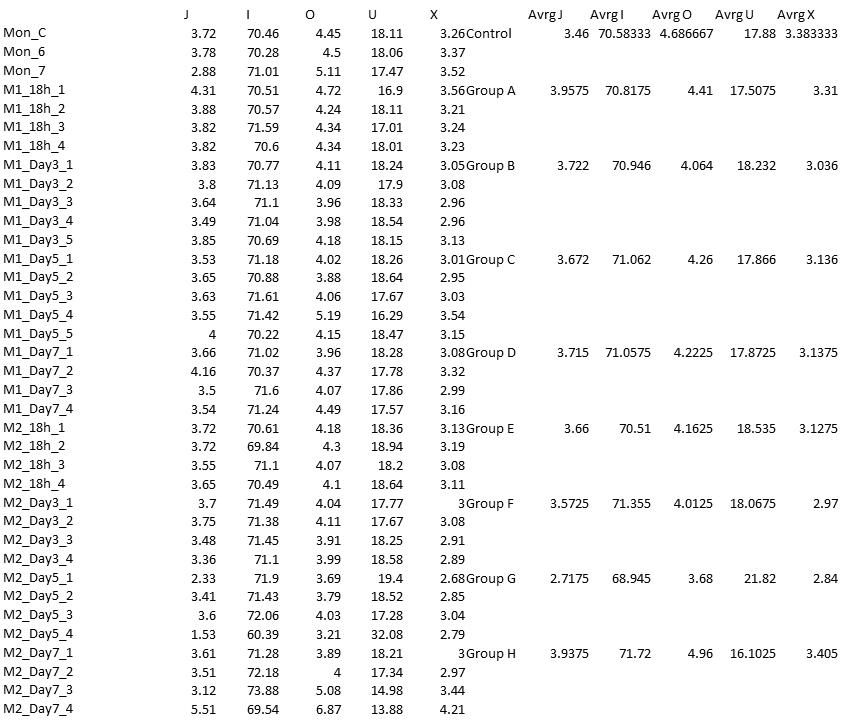

**Data S1.** Percentage of novel transcripts with class code i,j,o,u,x

**Data S2**. **Go_enrichment_lncRNA_M1 and M2 macrophages**. LncRNA paper. Supplementary figures and tables\Files\S2 file. GO_enrichment_lncRNA_M1 & M2 macrophages.xlsx

Data S3. LRRC75A-AS1 PCR array report. LncRNA paper. Supplementary figures and tables\Files\S4 file. LRRC75A-AS1 PCR array.xlsx

Data S4. GAPLinc PCR array report. LncRNA paper. Supplementary figures and tables\Files\S5 file. GAPLINC PCR array.xlsx
